# Deep learning for population size history inference: design, comparison and combination with approximate Bayesian computation

**DOI:** 10.1101/2020.01.20.910539

**Authors:** Théophile Sanchez, Jean Cury, Guillaume Charpiat, Flora Jay

## Abstract

For the past decades, simulation-based likelihood-free inference methods have enabled researchers to address numerous population genetics problems. As the richness and amount of simulated and real genetic data keep increasing, the field has a strong opportunity to tackle tasks that current methods hardly solve. However, high data dimensionality forces most methods to summarize large genomic datasets into a relatively small number of handcrafted features (summary statistics). Here we propose an alternative to summary statistics, based on the automatic extraction of relevant information using deep learning techniques. Specifically, we design artificial neural networks (ANNs) that take as input single nucleotide polymorphic sites (SNPs) found in individuals sampled from a single population and infer the past effective population size history. First, we provide guidelines to construct artificial neural networks that comply with the intrinsic properties of SNP data such as invariance to permutation of haplotypes, long scale interactions between SNPs and variable genomic length. Thanks to a Bayesian hyperparameter optimization procedure, we evaluate the performance of multiple networks and compare them to well established methods like Approximate Bayesian Computation (ABC). Even without the expert knowledge of summary statistics, our approach compares fairly well to an ABC based on handcrafted features. Furthermore we show that combining deep learning and ABC can improve performance while taking advantage of both frameworks. Finally, we apply our approach to reconstruct the effective population size history of cattle breed populations.

## 1 Introduction

In the past years, fields such as computer vision and natural language processing have shown impressive results thanks to the rise of deep learning methods. What makes these methods powerful is not fully understood yet, but one key element is their ability to handle and exploit high dimensional structured data. Therefore, deep learning seems particularly suited to extract relevant information from genomic data. It has indeed been used for many tasks outside population genetics, such as detection of alternative splicing sites, prediction of protein binding sites or other phenotype markers (Alipanahi et al., 2015, Jaganathan et al., 2019, Ma et al., 2018).

As genomic data becomes more and more available, it is now possible to leverage genetic variations within species or populations to investigate complex demographic histories including multiple admixture events, population structure or size fluctuation through time. In fact, initiatives like the 1000 Genomes Project for human populations (Consortium et al., 2010) have been extended for better world coverage and data quality (Bergström et al., 2019, Consortium et al., 2015, Leitsalu et al., 2014, Mallick et al., 2016, Pagani et al., 2016) and opened up to many other species such as *Bos taurus* with the 1000 Bull Genomes Project (Daetwyler et al., 2014) or chimpanzees and gorillas with the Great Apes Genome Project (Prado-Martinez et al., 2013). Even for smaller scale studies, researchers often have access to the whole genomes or high-density SNP data of numerous samples. These data collections can only be treated with inference methods able to scale to dozens or hundreds of individuals and large amount of genetic markers.

In this study, we propose several deep learning approaches for reconstructing the detailed histories of past effective population sizes from genetic polymorphism within a single population, a task considered difficult for various reasons. First, a present-day population, and even more so a sample of it, is one among many possible outcomes of a stochastic process depending on population sizes, mutations and recombinations. Second, many other factors such as selective pressure, admixture events or population structure also shape the contemporary genetic diversity, which can blur the link between population size history and genetic data. As a result, the accuracy of the reconstruction and its level of resolution depend on the number of individuals available, the quality of the data and the methodology used. Nonetheless, in practice previous methods such as Bayesian skyline plots and their derivatives (Ho and Shapiro, 2011), sequential Markov coalescent (SMC) (PSMC, diCal and their derivatives (Li and Durbin, 2011, Sheehan et al., 2013)), Approximate Bayesian Computation (Boitard et al., 2016b, Navascués et al., 2017) and SFS-based approaches (Bhaskar et al., 2015, Liu and Fu, 2015) have shown great results, supporting archaeological evidence and helping to understand species decline or expansion.

The study of genetic variation relies primarily on genotyping and sequencing data of very high dimensionality, which is a major difficulty for most inference methods. Some approaches, such as coalescent-HMMs methods (Spence et al., 2018), enable parameter inference using the full dataset by making simplifying assumptions on the underlying models. A few of them can process unphased data (Terhorst et al., 2017), scale to large sample size (Terhorst et al., 2017) or to complex models (Steinrücken et al., 2019). However, no method simultaneously addresses all three. Moreover, handling arbitrarily complex models remains untested (e.g. models with more than three populations) or intractable (e.g. complex spatial models) (Spence et al., 2018). Hence, most frameworks solving complex population genetic tasks do not rely on coalescent-HMMs and reduce the data dimension with a pre-processing step during which the dataset is converted into a smaller set of statistics called summary statistics. These statistics can then be used in likelihood and composite likelihood inference frameworks, when the model or statistics are simple enough, or in simulation-based approaches. Among the latter, the widely used Approximate Bayesian Computation (ABC) framework as well as several machine-learning algorithms, including Support Vector Machine (SVM) and random forests, were able to tackle a variety of tasks such as demographic model selection and parameter inference (Excoffier et al., 2013, Jay et al., 2019), detection of selection (Sugden et al., 2018, Tournebize et al., 2019) and introgression (Schrider et al., 2018). The current trend when addressing complex tasks is to include a large number of summary statistics inspired by population genetic theory in order to minimize the information loss. Summary statistics commonly used are the site frequency spectrum (SFS) and its summaries (e.g. Tajima D), linkage disequilibrium (LD) and statistics based on shared segments that are identical-by-state (IBS) or identical-by-descent (IBD) (Gladstein and Hammer, 2019, Jay et al., 2019, Sheehan and Song, 2016, Smith and Flaxman, 2019). However, they are not guaranteed to be sufficient and the inclusion of numerous statistics can impact the performance of standard ABC, a problem known as curse of dimensionality (Blum, 2010). An active research topic in the ABC community is thus the development of methods addressing this curse of dimensionality by (i) selecting the best subset of summary statistics according to some information-based criteria, or (ii) integrating machine learning steps into ABC to handle a larger number of summary statistics (e.g. kernel methods, random forests), or (iii) constructing summary statistics using linear and non-linear models based on candidate statistics or on the original data when feasible (Aeschbacher et al., 2012, Blum et al., 2013, Fearnhead and Prangle, 2012, Jiang et al., 2017, Nakagome et al., 2013, Raynal et al., 2018).

In our study, we use deep learning, a method derived from machine learning. The objective of this method is to design a function, represented by an artificial neural network (ANN), which is a differentiable computational graph organized as a stack of linear and non-linear layers, with a high number of trainable parameters (usually thousands or millions). A network layer takes as input the outputs of the previous layer(s): each node of the layer performs a linear combination of the inputs, followed by a non-linear transformation, and this value is passed to the next layer. Networks vary in their shape (number of layers and nodes) and in the way nodes are connected. For example a Multi Layer Perceptron (MLP) connects all nodes of a layer to all nodes of the following layer (Rumelhart et al., 1986), while a Convolutional Neural Network (CNN) connects only nodes of similar location (LeCun et al., 1995). Despite the differences, any network defines a parameterized function that allows for a complex non-linear mapping from a space to another, and therefore can solve a complex task, when the provided parameters are suitably adjusted. To tune the parameters, the network is trained, thanks to a *training set* consisting of examples of (input, desired output) pairs, by optimizing a criterion (*loss function*), that expresses how well the network performs on the dataset with its current parameters. For example, for an object recognition task in images, the input is an image, the output is a probability distribution over possible names of objects, and the loss is the distance between the prediction of the ANN and the expected output (a Dirac peak on the name of the object shown by the image). The parameters of the function are tuned to minimize this loss thanks to an optimization algorithm based on gradient descent and backpropagation. This process usually requires a large training dataset, in order for the network to be able to learn and generalize well, that is, to perform well on data never seen so far.

Deep learning has only recently been used to tackle population genetics questions. First, multilayer perceptron (MLP) were used to process small SNP windows for population assignment (Bridges et al., 2011). Then, the same type of architecture has been used to process large sets of summary statistics for predicting jointly selective sweeps and simple demographic changes (Sheehan and Song, 2016). Villanea and Schraiber (2019) also applied MLP on summary statistics to discriminate between multiple scenarios of archaic introgression and two other studies added an ABC step to address a similar task (Lorente-Galdos et al., 2019, Mondal et al., 2019). A second type of ANN, convolutional neural networks (CNN), were then applied to summary statistics computed over 5Kb genomic regions in order to predict selective sweeps (Xue et al., 2019). A considerable shift occurred when several studies applied ANN directly on genomic data instead of using summary statistic. Various CNN architectures processing SNP matrices were proposed to infer recombination rates along the genome (Chan et al., 2018, Flagel et al., 2018), selection (Flagel et al., 2018, Torada et al., 2019), introgression (Flagel et al., 2018) and three-step population size histories (Flagel et al., 2018). The CNN implemented by Chan et al. (2018) and based on Deep Sets (Zaheer et al., 2017) is invariant to haplotype (chromosome) permutation, i.e to the permutation of rows in the SNP matrix, thanks to convolution filters that treat each haplotype in an identical way. The other approaches proposed instead to sort haplotypes by similarity before processing them with filters sensitive to the haplotype order (Flagel et al., 2018, Torada et al., 2019). More recently, Recurrent Neural Networks (RNN) were applied to estimate the recombination rate along the genome (Adrion et al., 2019), and Generative Adversarial Networks (GAN) to learn the distribution of genomic datasets and generate artificial genomes (Yelmen et al., 2019).

Convolution layers have been particularly efficient for large size data with spatial coherence such as images, exploiting the geometric structure of the image pixel grid. Instead of requiring as many weights as the input data size (like fully-connected layers in MLP), convolution layers take advantage of the spatial structure of the data, by defining spatially-small filters and applying them at each location along the input dimension (here, the SNP sequence). The result of each operation is ordered spatially according to the corresponding location in the input, to keep spatial coherence throughout the network. The filters of the first convolution layers have a small scope over the data input of the network, but adding layers on top of each other gives a larger scope to the last layers, allowing to handle long range interactions, in particular when combined with max-pooling layers. These interactions are important for demographic inference, e.g. they can allow the network to measure linkage disequilibrium and thus, the level of recombination.

Among the variety of developed ANN architectures, it is not straightforward to know which one is the most adapted to genomic data for a given population genetic task. In particular, this study aims at reconstructing detailed step-wise effective population size histories with 21 size parameters under an unknown recombination rate, a complex model with a fairly high dimensional parameter space compared to the population genetic task previously addressed with ANN. Hence we propose multiple networks, some of which are new and designed specifically for population genomics, and others are more basic. We then apply a hyperparameter optimisation procedure (BOHB (Falkner et al., 2018)) to select the best architecture and hyperparameters. We investigate the performance of two MLPs, one using summary statistics and one using SNPs data of fixed length. We also compare two novel CNN based architectures, one with mixed convolution filter sizes over multiple individuals and another CNN that is adaptive to the genomic input size and invariant to the permutation of individuals or haplotypes. Both networks incorporate SNP data and their positions (encoded as distances between SNPs), a concept also developed in a different fashion by Flagel et al. (2018). In our last setup, we combine ABC and ANN by using the ANN predictions as summary statistics with the aim to benefit from both method advantages. Because no end-to-end deep learning approach for demographic inference had yet been compared to ABC or other traditional methods, we carefully benchmarked all these networks against variations of PopSizeABC, one of the highly performing methods for step-wise size inference that is based on ABC (Boitard et al., 2016b). We also compare our architecture with CNNs developed for a related demographic task (Flagel et al., 2018). Finally we apply our approach to real genomes in order to reconstruct the size history of three cattle breeds.

## 2 Materials and Methods

In this study, we introduce the first deep learning approaches for inferring detailed histories of effective population sizes using genomic data. Based on whole sequences of SNP data of multiple individuals from a single population, we aimed to predict 21 population size parameters, each corresponding to a time step. Our method and the baseline frameworks all relied on large-scale simulated datasets for which the true demographic parameters are known and drawn from prior distributions of population sizes and recombination rates. For each drawn parameter set (i.e. demographic scenario), we simulated 100 independent genomic loci of length 2Mb (i.e. 100 replicates) for 50 haploid individuals using msprime (Kelleher et al., 2016). Using this reference panel, we then trained methods based on ABC, deep learning or a combination of both, to predict the demographic parameters (Figure 1). In this section, we will give an overview of these methods as well as the hyperparameter optimisation procedure.

**Figure 1:**
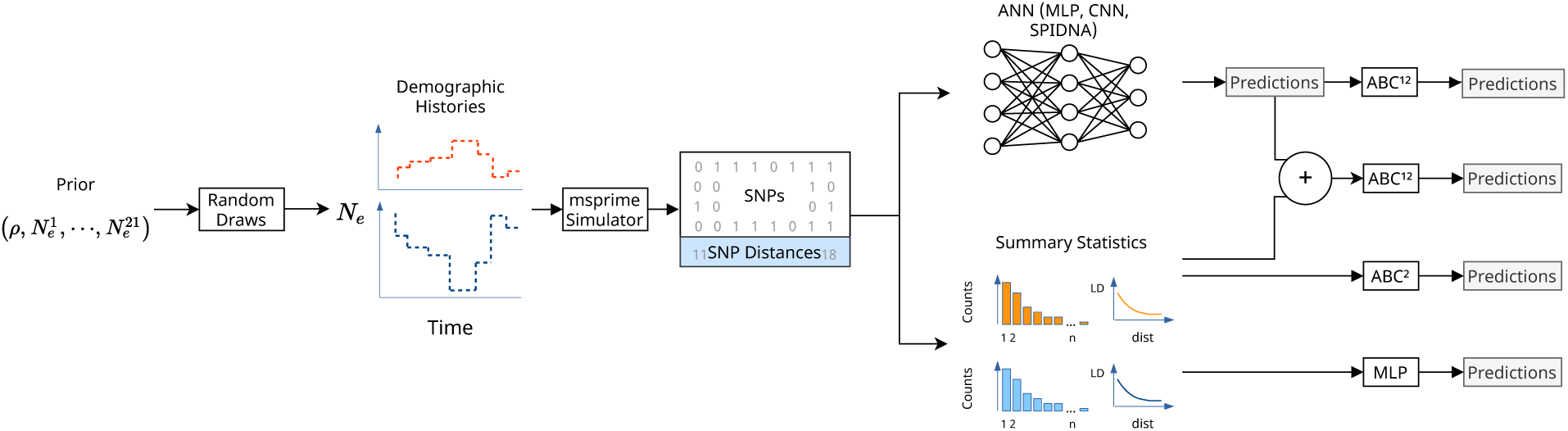
Overview of the methods compared in this study. Demographic histories are drawn from a prior distribution on 21 population sizes 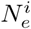 and one recombination rate *ρ*, and are used to generate SNP matrices with msprime. Two types of summary statistics are computed from these simulations (SFS and LD). The predictions (outputs) made by different kind of ANNs (MLP, *custom* CNN and SPIDNA architecture) are compared to an MLP using the summary statistics and to ABC using either the summary statistics, SPIDNA outputs or both. 1 ANN outputs used are the predictions made by the version of SPIDNA with the lowest prediction error. 2 ABC without correction, with linear regression, ridge regression or a single layer neural network are compared.

### 2.1 Simulated data and summary statistics

#### Neutral simulations

All methods compared in this study are trained in a supervised fashion, and thus require genetic data of numerous populations under various demographic scenarios. We defined the demographic parameters by following similar rules as Boitard et al. (2016b): *I* = 21 time windows [*t*_*i*_, *t*_*i*+1_] were defined from present to ancient periods with 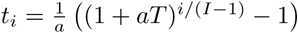 generations, *i* going from 0 to *I* −1, *T* = 130, 000, *a* = 0.06 and *t*_*I*_ = +*∞*. These values of *T* and *a* were chosen by Boitard et al. (2016b) to capture important periods of cattle history. They could be modified to describe more precisely specific parts of the history by playing with the ratio between the length of recent versus old time windows. By increasing exponentially the time windows as we go further in the past, we obtain more detailed scenarios for recent times. The time windows are identical for all scenarios. Each demographic scenario is generated by drawing a first population size *N*_0_ between 10 and 100,000 from a uniform distribution and corresponds to the most recent time window *t*_0_. The population sizes of the next time windows follow *N*_*i*_ = *N*_*i*−1_ × 10^*β*^ with *β* sampled uniformly between −1 and 1. *β* is redrawn if it gives a population size out of]10; 100, 000[. We randomly drew from this prior distribution 50,000 scenarios and simulated 100 independent 2Mb-long segments of 50 haploid individuals for each scenario, using the msprime coalescent simulator version 0.6.1 (Kelleher et al., 2016). We obtained a total of 5,000,000 SNP matrices *X* of size *M* = 50 haplotypes × *S* SNP sites, each associated to a vector of size *S* that contains the distances between SNPs (in bp). Ancestral and derived alleles are encoded with 0 and 1. The mutation rate is set to 10^*−*8^ as in MacLeod et al. (2013). The recombination rate is sampled uniformly between 10^*−*9^ and 10^*−*8^ for each scenario to be consistent with the estimations in cattle breeds (Sandor et al., 2012). In order to compare all methods based on the same training panel we set a minimum threshold of 400 SNPs per 2Mb region and designed the networks accordingly. Scenarios producing less than 400 SNPs in any 2Mb regions were removed. This threshold could be changed by modifying the networks or simulating longer regions. However, the real cattle dataset has on average 4,357 SNPs across a 2Mb-long region, so these scenarios were far outside the plausible posterior distribution. That reduced the dataset to 18,461 scenarios (i.e. 1,846,100 SNP matrices) out of the 50,000 scenarios simulated with an average of 2,486 SNPs and a maximum of 17,839 SNPs. This dataset is split into a validation set of 500 scenarios (i.e. 50,000 validation SNP matrices overall) and a training set with the remaining 17,961 scenarios (i.e. 1,796,100 training SNP matrices). In order to check for hyperparameter overfitting, we have also simulated a test set from the same prior distribution. Hence, we randomly drew 2,000 scenarios and kept the 767 scenarios with more than 400 SNPs which gives 76,700 test SNP matrices. Training, validation and test set demographic parameters were all standardized using mean and variance from the training set.

Except stated otherwise, methods that are not adaptive to the number of SNPs used only the first 400 SNPs of each SNP matrix. The proportion of these 400 SNPs kept among all SNPs from a simulated matrix is on average 28%.

We make the assumption that the 2Mb-long windows of a scenario are independent, which is true for simulated data but not for real data. Information across windows (100 windows by scenario for simulated data and 1,213 for real data) is combined during the summary statistics computation step for methods using summary statistics or by averaging the network predictions over all windows for methods using SNP matrices as input. Thus, the spatial information that may exist across these windows for real data is not conserved.

#### Simulations with selection

To investigate the robustness of our approach, an extra set of data was simulated under demographic changes and selective pressure. We used *msms* (Ewing and Hermisson, 2010) to simulate scenarios including positive selection with additive fitness using varying values of selection coefficient (*s* in 2Ne units: 100, 200, 400 or 800), selection starting time (*T*_*sel*_: 200, 1000 or 2000 generations ago) and initial frequency of the beneficial allele (*f*_0_: 0.1%, 1%, 5%). The SNP under selection was located at the center of the region. The mutation rate was set to 10^*−*8^, the recombination rate to 5 × 10^*−*9^, the number of haplotypes to 50 and the region length to 2Mb. We generated 16 × 100 replicates for each of the 36 selection parameter combinations (*s*, *T*_*sel*_, *f*_0_) and 30 × 100 replicates with no selection under three demographic scenarios (constant, declining or expanding size) leading to a total of 181,800 SNP matrices. Inference methods requiring a fixed input size processed the 400 successive central SNPs (ie 200 before and 200 after SNP under selection).

#### Summary statistics

For each group of 100 segments corresponding to one scenario, we computed the site frequency spectrum and the linkage disequilibrium as a function of the distance between SNPs averaged over 19 distance bins for a total of 68 summary statistics. Our python script is partly based on the scikit-allel python module (Miles et al., 2019). These predefined summary statistics constitute the training, validation and test set for all methods based on summary statistics or on their combination with SNP matrices.

### 2.2 Baselines

We compared our approach to five baselines: an ABC approach and a MLP both using linkage disequilibrium and site frequency spectrum as summary statistics, and another MLP, a *custom* CNN and a CNN from (Flagel et al., 2018), all using genomic data directly. We evaluated four ABC methods (rejection, local linear regression, local ridge regression and a single-hidden-layer neural network). Their performance represents a baseline of commonly used methods to be compared to ours. The second framework, a MLP processing the same summary statistics as ABC, is similar in spirit to the one previously proposed for predicting three demographic parameters alongside selection (Sheehan and Song, 2016). It is the baseline for methods using deep learning based on summary statistics. The third and fourth baselines consist in a MLP and a CNN processing directly the first 400 SNPs of a 2Mb-long genomic region instead of summary statistics. The MLP takes as input SNP data and locations flattened into a single vector. In this configuration, the spatial information between SNPs, the link between the SNPs and their location, and the link between alleles of the same individual are lost. The fourth baseline is a newly design CNN with rectangular shaped filters that cover more than one haplotype (i.e. more than one row). It takes as input a matrix in which rows are the haploid individuals and columns are the genetic markers. An additional row contains the SNP positions encoded as the distances between SNPs. The input size is (number of haplotypes+1) × number of kept SNPs. This *custom* CNN is a first step towards an architecture tailored to raw genomic data, because spatial information is preserved as for recent ANNs applied to population genetics (Chan et al., 2018, Flagel et al., 2018, Torada et al., 2019), but also because asymmetrical filters of various sizes account for the heterogeneous entities of axes (haplotype versus SNP, rather than pixel versus pixel). Finally we adapted and re-trained four networks among the top-ranked CNNs proposed by Flagel et al. (2018) so that they could reconstruct a 21-epoch model of instantaneous effective population size rather than the three-epoch model initially investigated by the authors, and for practicability we called them *Flagel* CNNs.

Each method is evaluated using its prediction error given by the following mean squared error:

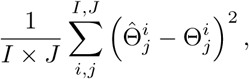

where 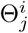 and 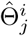 are respectively the true and predicted standardized population size for the time window *i* and scenario *j*, *I* = 21 is the number of time windows and *J* the number of scenarios in the set. For inference based on raw data and neural networks, the prediction 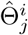 is given by the average of the population sizes 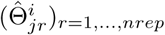 estimated for each replicate (independent region) *r*.

#### Approximate Bayesian Computation

We compared ABC with the simple rejection procedure (i.e. no correction) and three correction methods implemented in the R package ‘abc’ (Csilléry et al., 2012): local linear regression, ridge regression and non-linear regression based on a single-hidden-layer neural network. Settings were set to default except for the tolerance rate set to six possible values (0.05, 0.1, 0.15, 0.2, 0.25 and 0.3). ABC was run on (a) predefined summary statistics, (b) SPIDNA outputs (i.e. automatically computed summary statistics), or (c) a combination of predefined summary statistics and SPIDNA outputs. We used the median of the posterior distribution as the demographic parameter estimate 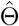.

#### Multi-Layer-Perceptron Networks

The first MLP is based on summary statistics, has 3 hidden layers, ReLU activation functions and uses batch normalisation. As in Sheehan and Song (2016), the hidden layers have respectively 25, 25, and 10 neurons. It takes 35 summary statistics as input. This network and all the following ones output 21 demographic parameters and are trained with a regular L2 loss function and adam optimizer (Kingma and Ba, 2014) unless stated otherwise. This MLP has a total of 2,986 trainable parameters. Our second MLP is based on ‘raw’ genomic data and takes a matrix of 50 haplotypes (rows) for 400 SNPs (columns) and its associated vector of distances between SNPs as input. Its hidden layers respectively have 20, 20, and 10 neurons, which gives it 408,981 trainable parameters.

#### *Custom* CNN

Our convolutional neural network takes as input the same matrix of 400 SNPs and has 2-dimension filters of various shapes. The first layer consists of 5 kernels with rectangular shape (2×2, 5×4, 3×8, 2×10, 20×1) applied to the SNP matrix *X*. Each kernel creates 50 filters, which amounts to 250 feature maps after the first layer. The SNP distance vector *d* is treated by the 5 associated kernel shapes (1×2, 1×4, 1×8, 1×10, 1×1) with 20 filters each, making 100 filters in total. The results of the first convolutional layer are then concatenated so that the second convolutional layer will couple information from *X* and *d* in a way that emphasizes the original location of the SNPs along the genome. The outputs of this second layer are then combined and go through 5 convolutional layers and 2 fully connected layers. Adding convolutional layers one after the other allows our network to combine patterns and reduce the size of the data without adding too many weights to our model. This network has a total of 131,731 trainable parameters.

#### *Flagel* network

We reused the code associated with the repository of the first paper using a CNN for demographic inference (Flagel et al., 2018) and adapted it to our dataset and task. We trained the network with the exact same architecture as the one published, except that we changed the last layer to allow the prediction of our 21 population size parameters. We parametrized the network with the set of hyperparameters leading to the best performance in the previous work for two different types of SNP encoding (0/255 or −1/1). It is noteworthy that the actual encoding in their code is 0/-1 and not 0/255, thus we kept the same encoding to be able to compare the performance. The networks were trained with the same procedure of 10 epochs with early stopping in case of no progression of the loss after 3 epochs. The batch size is 200. The input data had 50 haplotypes and either 400 SNPs as processed by our *custom* CNN or we downsampled the data to one every ten SNPs as done in the original work, leading to 1,784 wide input SNP matrices. This size corresponds to the tenth of the biggest SNP matrix in our dataset. Smaller simulations are padded with zeros. All parameters can be found in table S1.

### 2.3 Sequence Position Informed Deep Neural Architecture

We called our architecture SPIDNA, for Sequence Position Informed Deep Neural Architecture, and designed it to comply to the principal features of SNP data: data heterogeneity (data includes genetic markers and their positions encoded as distances between SNPs), haplotype permutation invariance, long range dependencies between SNPs and variable number of SNPs. Similarly to our *custom* CNN, SPIDNA takes as input a matrix describing haploid individuals as rows and SNP as columns, with an additional row for the SNP distances.

#### 2.3.1 Permutation invariance

One of the SNP matrix properties is its invariance to the permutation of haploid or diploid individuals (rows of the SNP matrix). The same matrix with permuted rows contain the exact same information and should lead to the same predictions. Most summary statistics are already invariant to the haplotype order by definition. On the other hand, typical operations used in ANN such as rectangular filters and fully connected layers are not invariant, and consequently our two baseline ANNs do not respect this data feature. Here we implemented an architecture invariant by design, that stacks functions equivariant and invariant to row permutations (Lucas et al., 2018). It is a modification of the Deep Sets scheme (Zaheer et al., 2017) used for population genetics under the name “exchangeable networks” (Chan et al., 2018). In our study, the equivariant function is a convolutional layer with filters of size 1 × *a*, that treats each haplotype (row) independently and computes equivariant features, while the invariant function computes the mean of these features over the row dimension. The invariant function reduces the dimension of the data to one row, which is then concatenated to each equivariant row (Figure 2). Therefore the correlation between rows increases at each layer, which progressively transforms the equivariant input to an invariant output. However, the correlation increase should be moderate and progressive to avoid immediate loss of the information at the haplotype level. To promote this, we perform two independent normalizations, one over the output of the equivariant function and one over the input of the invariant function, and associate a correlation control parameter *α* that quantifies the contribution of the invariant function to the next layer, thus controlling the speed at which the correlation increases between rows.

**Figure 2:**
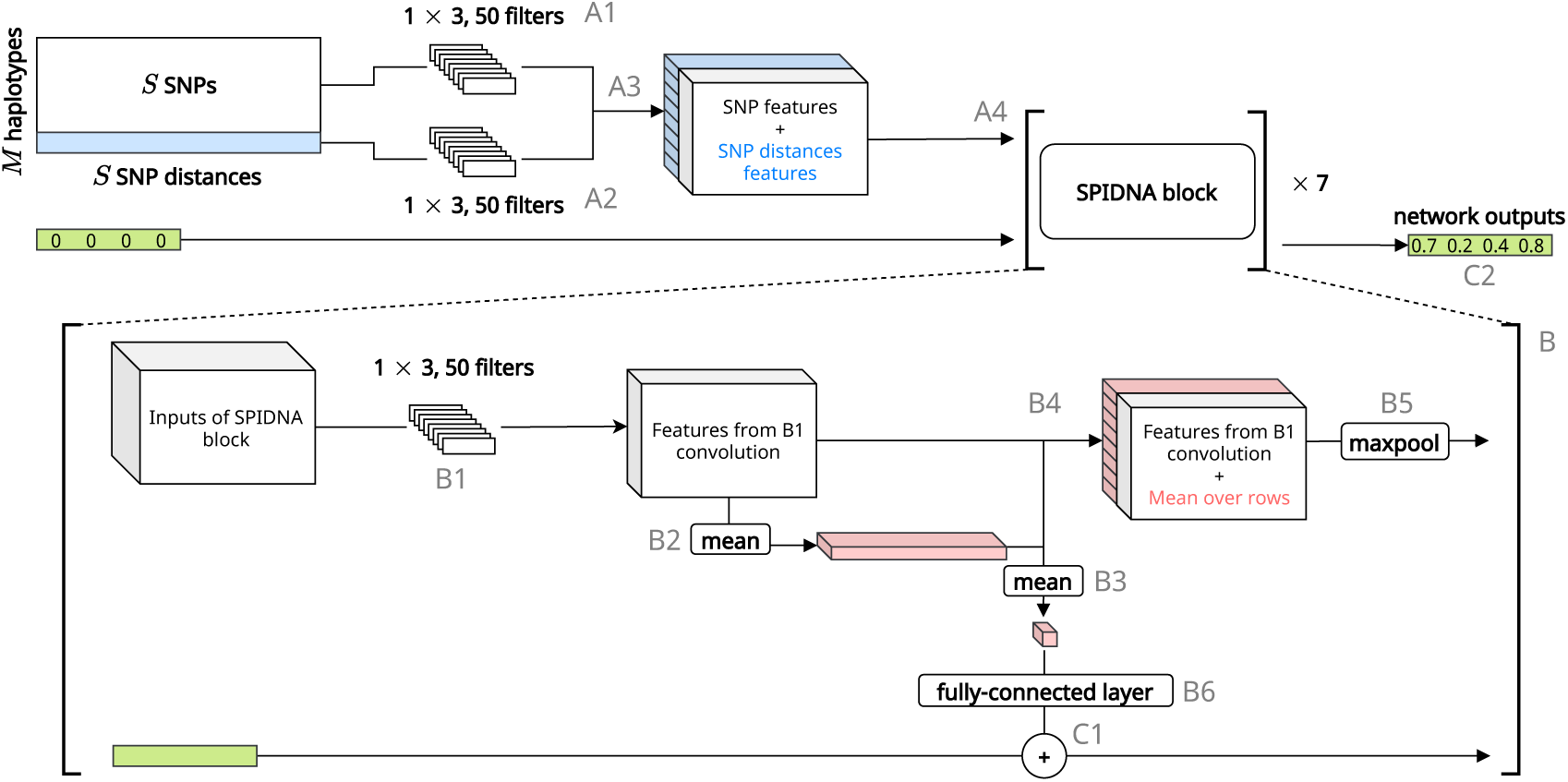
Schematic of SPIDNA architecture. SPIDNA takes as input a SNP matrix associated with its vector of distances between SNPs (in blue). A convolution layer is applied to the SNPs (A1) and another convolution layer is applied to the distances (A2). Results of A2 are repeated to be concatenated with results from A1 (A3). The output is passed to a series of seven SPIDNA blocks (A4). Each SPIDNA block starts with a convolution layer (B1) followed by the mean over rows of the convolution layer result (B2) and the mean over columns of B2 result (B3). The concatenation of B1 and B2 results (B4) is processed by a max pooling layer (B5) and passed to the next SPIDNA block. In parallel, the output of B3 is processed by a fully-connected layer (B6). The prediction vector (in green) is updated at each SPIDNA block with a sum (C1) of its previous value and B6 results. It is finally output by the last block as the predicted demographic parameters (C2).

#### 2.3.2 Convolution networks to handle data with variable size

A major difficulty that arises with genomic data is that the number of SNP varies from one dataset to another, or from one genomic region to another, due to the stochasticity of biological and demographic processes (and of their corresponding genetic simulations). Therefore, we use convolution layers as they can handle data with variable size while keeping the number of network weights constant. A filter can be repeated more or fewer times to cover the whole input entering each layer, letting the network adapts itself to the data. Consequently, the output size of each convolution layer will vary depending on the input size. This prevents the use of fully connected layers directly after a convolution layer as it is often the case with CNNs. Instead, we use fully-connected layers only after operations independent of the input size and with a fixed output size, namely mean functions over the column and row dimensions (Figure 2).

Overall, we designed an architecture accounting for invariance and adaptive specificities by stacking multiple equivariant blocks (Figure 2, label B). An equivariant block consists in one convolution layer with filters of size 1 × 3 that are equivariant (B1), averages of the convolution outputs across the haplotype axis (B2) and the SNP axis (B3) that are both invariant, a concatenation of the equivariant and invariant features (B4), one max pooling layer that is also adaptive to the number of SNPs (B5) and one fully-connected layer that updates the demographic predictions at each block (B6) via a sum function (Figure 2). Contrary to the MLPs, *custom* CNN and non-adaptive SPIDNA that include batch normalization of neuron activities, adaptive SPIDNA networks include instance normalization.

For our permutation invariant architecture, we designed three variations. The first one uses batch normalization, after each convolution layer, and therefore takes as input a fixed number of 400 SNPs, similarly to two of the baselines. The second one is invariant to the number of SNPs and uses instance normalization, after each convolution layer, to normalize layer inputs per-data instead of per-batch (for the batch normalization). The last variation is also invariant to the number of SNPs, but uses two separate instance normalization steps, as well as the correlation control parameter *α*.

Except for these differences, these three variations have the same architecture, represented in Figure 2. At each step *i* of the network, we consider that the data has four dimensions *B*_*i*_ × *M*_*i*_ × *S*_*i*_ × *F*_*i*_, *B* being the batch dimension, *M* the row dimension (also the haplotype/genotype dimension before the first layer), *S* the column dimension (also the SNP dimension before the first layer) and *F* the feature dimension (only one feature before the first layer). A first convolution layer of 50 1×3 filters is applied to the SNP matrix (Figure 2, label A1), and another convolution layer of 50 1×3 filters is applied to the vector of distances between SNPs (A2) and repeated *M* times. The results of the two convolutions have now the same dimensions and are concatenated along the feature dimension (A3). The resulting tensor is then passed to seven blocks put end to end (A4), each one involving an equivariant function and an invariant function (B). The equivariant function *ψ* is a convolutional layer of 50 1×3 filters (B1) that outputs a tensor of size *B*_*i*−1_ *× M*_*i*−1_ × (*S*_*i*−1_ *−* 2) *× F*_*i*−1_/2. The result of the equivariant function is then passed to the invariant function *ρ*, which is the mean over the dimension *M* (B2). Thus *ρ*(*φ*(*X*_*i*−1_)) has size *B*_*i*−1_ × (*S*_*i*−1_ *−* 2) *× F*_*i*−1/2_, which is repeated *M* times to maintain the same dimension as *φ*(*X*_*i*−1_). Then *ρ*(*φ*(*X*_*i*−1_)) and *φ*(*X*_*i*−1_) are concatenated over the feature dimension (B4). Finally, max-pooling filters of dimension 1×2 are applied, and the result is passed to the next block (B5). In parallel, each block computes the average over the column dimension *S* of the 21 first features of *ρ*(*φ*(*X*_*i*−1_)) that are then passed to a fully-connected layer with 21 outputs (B6). The predictions of each block are summed (C).

### 2.4 From batch normalization to instance normalization

Network weight initialization is a difficult task that can lead to vanishing or exploding gradient when not carefully done and associated with a poor learning rate (Bengio et al., 1994, Glorot and Bengio, 2010). Most initialization schemes try to force the outputs of each layer to follow some distribution assuming normalized input data. Batch normalization solves this problem by normalizing layer outputs over the whole batch during training and computing a running mean and variance for the evaluation steps. We used this type of normalization for our networks that take as input a fixed number of SNPs. For the networks invariant to the number of SNPs, we could not concatenate all batch data into the same tensor because of their varying sizes. Therefore, we use instance normalization, which computes both mean and variance over the feature dimension.

### 2.5 Hyperparameter optimization

Compared to other machine learning methods, ANNs have a potentially infinite amount of hyperparameters when including for instance the number of layers, the number of neurons in each of them, the learning rate, weight decay or the batch size. Moreover, a run over a full dataset with enough epochs to reach convergence is time consuming for networks with a complex architecture defined by many learnable parameters. Therefore, the development of deep learning architectures often relies on the experience and intuition of the practitioner in a try-and-repeat process. Grid search and random search are two strategies for exploring the hyperparameter space uniformly. They are commonly used but are limited by the computing resources available. In our study, we used HpBandSter, a package that implements the HyperBand (Li et al., 2016) algorithm to run many hyperparameter trials on a smaller resource budget (i.e. few epochs) and runs the most promising trials on a greater budget. Combined with BOHB (Falkner et al., 2018), a Bayesian optimisation procedure that models the expected improvement of the joint hyperparameters, this method provides more guided and faster search of the hyperparameter space. At each step, BOHB draws a new combination of hyperparameter values to be tested according to the expected improvement and to a predefined prior. Here, we performed a search in a 5-dimensional space defined by uniform priors over the type of architecture (architectures from our baselines and variations of SPIDNA architecture, based on 400 SNPs or the full number of SNPs), the learning rate, the weight decay and the batch size. For SPIDNA architectures that controlled correlation, we added the control parameter *α* to the Bayesian optimization procedure with a log-uniform prior between 0.5 and 1. The search was performed for 3 budget steps and replicated 5 times, leading to a total of 83 successfully trained networks.

For the ABC baselines, the tolerance rates ranged from 0.05 to 0.3 by step of size 0.05 and were optimized for 12 ABC algorithms independently (4 correction methods × 3 types of inputs: predefined summary statistics, SPIDNA outputs or both).

As the training time of the MLP using summary statistics was short, we optimized its hyperparameters with a random search by drawing 27 configurations from uniform distributions and trained a network for each configuration during 6 epochs. The batch size was drawn between 10 and 100, learning rate between 5 · 10^*−*5^ and 1 · 10^*−*2^ and weight decay between 5 · 10^*−*5^ and 1 · 10^*−*2^.

### 2.6 Cattle breed data

We inferred the demographic history of Angus, Fleckvieh and Holstein cattle breeds using the data set of 25 sequenced individuals from the 1,000 genome bull project (Daetwyler et al., 2014) that was analysed by (Boitard et al., 2016b). As the data of real cattle sequence is prone to phasing and sequencing errors, we converted the real data from haplotype to genotype with a minimum allele frequency (maf) of 0.2, as suggested by Boitard et al. (2016b) and applied the same treatment to the simulated training set. We split the real data of each breed into 2Mb and removed segments comprising centromeres leaving 1,213 segments. We obtained a similar number of SNPs for the three breeds: Angus (average: 4,536 SNPs, maximum: 22,391 and minimum: 775), Fleckvieh (average: 4,837 SNPs, maximum: 24,896 and minimum: 896) and Fleckvieh (average: 4,732 SNPs, maximum, 24,098 and minimum: 1,212). Then we trained ABC, SPIDNA and a combination of both with the best hyperparameter configurations on the modified simulated data and performed the inference. The best version of SPIDNA without ABC is non-adaptive and therefore uses 400 SNPs from each segment which represents 10% of the total number of SNPs in the cattle data and 67% for the training dataset.

All computational resources used for this study are described in the Supplementary Text.

## 3 Results

### 3.1 Hyperparameter optimization

The configuration with the lowest loss generated by the hyperparameter optimization procedure used 400 SNPs with SPIDNA, batch normalization, a weight decay of 2.069 · 10^*−*2^, a learning rate of 1.416 · 10^*−*2^ and a batch size of 78 (Figure S1). Configurations with large batch sizes tended to have lower losses (Figure S1), which is expected as large batches provide a better approximation of the full training set gradient. However, a batch size too close to the training set size can lead to overfitting the training set. Here, we did not observe overfitting for any run when monitoring training and validation losses. The best configurations also tended to have low learning rates and weight decays (Figure S1). These low values slow down the convergence, but usually decrease the final prediction error if the budget (i.e. number of training epochs) is high enough for the network to reach convergence.

### 3.2 Comparison of the optimized architectures

For each architecture, we selected the best configuration obtained with the hyperparameter optimization procedure and trained it for a greater budget (i.e. 10 epochs), allowing an in-depth comparison. We found no strong decrease of prediction errors after this longer training compared to their counterparts with a 10^7^ budget (10^7^ training SNP matrix, i.e. 5.57 epochs) (Figures 3 and S1). Predictions errors for the validation set (used in the hyperparameter optimization procedure) and the test set are shown in the table S2. In the following paragraph, each method is designated along its index in the table.

**Figure 3:**
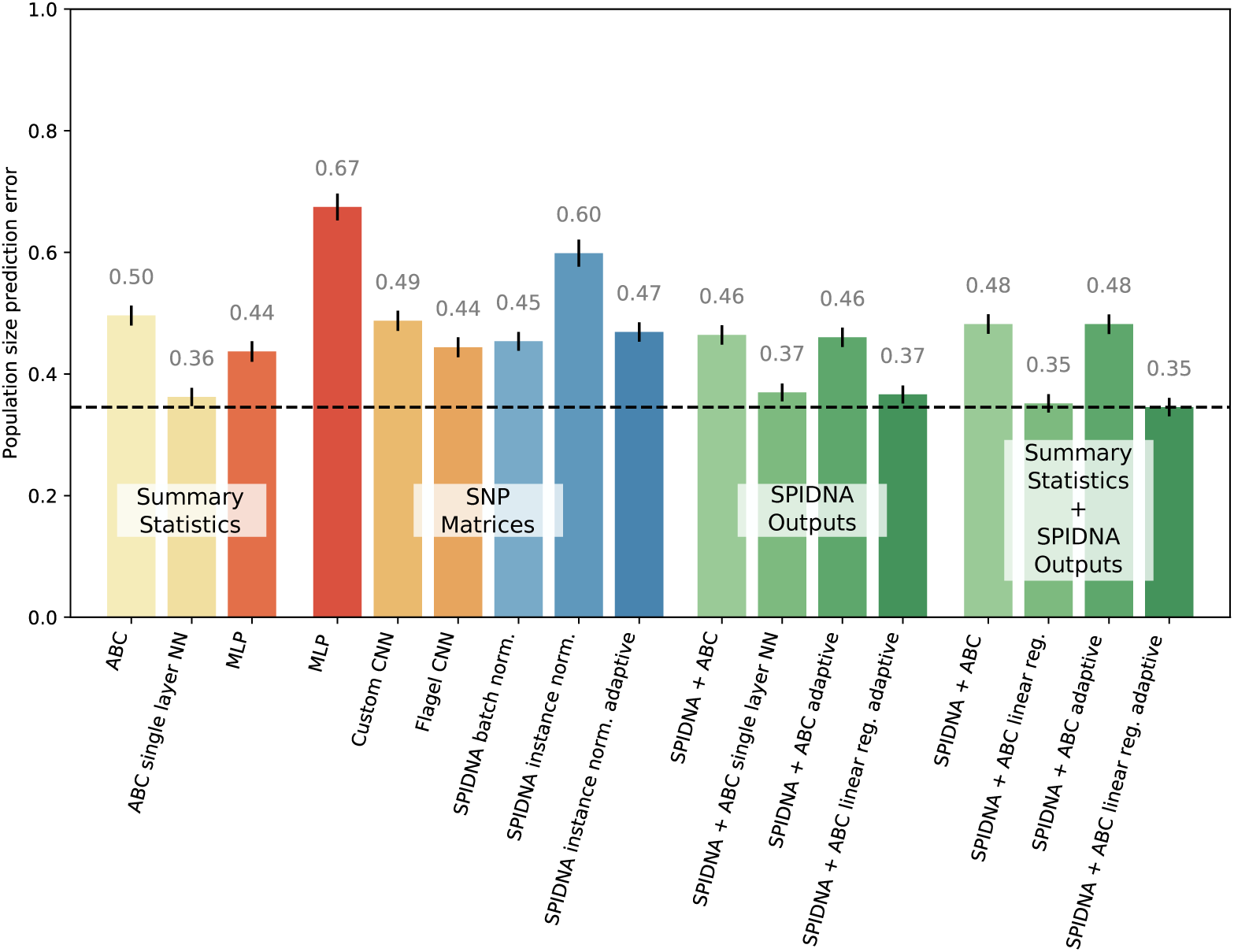
Prediction errors on the test set of the best run of each method after the hyperparameter optimization. The best configurations of each ANN (MLP, *custom* CNN and SPIDNA) have been retrained for 10 epochs. Traditional ABC methods are depicted in yellow, deep MLPs and CNNs in red and orange, SPIDNA ANNs in blue, combinations of ANNs and ABC in green. Methods are grouped into 4 families: “Summary statistics” (processed by ABC or ANN), “SNP matrices” (processed by ANN), “SPIDNA outputs” (processed by ABC, no summary statistic used), “Summary statistics and SPIDNA outputs” (processed by ABC). Vertical black lines on top of each bar represent the 95% confidence interval of prediction errors. Horizontal dashed line indicate the lowest error obtained (adaptive SPIDNA + ABC with local linear regression using summary statistics and SPIDNA outputs).

We first compared the optimized neural networks to optimized ABC approaches based on predefined summary statistics. The prediction errors achieved by ABC using summary statistics ranged from 0.496 (index 0, ABC rejection, i.e. without correction) to 0.364 (ABC neural networks, index 3). The MLP network based on summary statistics performed worse than ABC with correction (0.437, index 4). Moreover, MLP based on raw data performed very poorly (0.675, index 5) and all other networks based on raw data outperformed this MLP. Most of them (all except SPIDNA instance normalization on 400 SNPs, 0.641 and 0.599, index 12 and 14) outperformed the ABC rejection (0.454 and 0.469, index 11 and 15) or led to similar errors (0.489, index 13). The *Flagel* CNNs adapted from Flagel et al. (2018) that were not using dropout had average test losses of 0.541 and 0.444 (index 7 and 8). The two other *Flagel* networks achieved prediction errors similar to SPIDNA (network based on the first 400 SNPs: 0.609, index 9; network based on 1784 downsampled SNPs: 0.484, index 10), however they had 8 to 34 times more parameters than SPIDNA. Lastly, we evaluated two methods that combine deep learning and ABC, by considering the features automatically computed by a network as summary statistics for ABC (Jiang et al., 2017). When using only the predictions of SPIDNA as input to ABC with correction (linear regression, ridge regression or neural network), we improved greatly SPIDNA’s performance and obtained errors similar to the ABC based on predefined summary statistics (0.369 compared to 0.364, index 21 and 3). When using both SPIDNA predictions and predefined summary statistics as input to the ABC algorithm we decreased further the prediction errors (0.347, index 29).

### 3.3 Reconstruction of specific demographic histories using SPIDNA and SPIDNA+ABC

We further illustrated the performance of SPIDNA on a subset of demographic scenarios that were previously investigated (Boitard et al., 2016b) (Figure 4). We simulated six scenarios: “Medium”, “Large”, “Decline”, “Expansion”, “Bottleneck” and “Zigzag” the same way as the neutral simulations by specifying the demographic parameters instead of drawing them from a prior. The method correctly reconstructed histories of constant size, expansion and decline, as SPIDNA predictions from 100 independent genomic regions (black boxplots) approximately followed the real population size trend and magnitude. The true parameters were always included in the 90% credible intervals (light blue envelopes) predicted by SPIDNA combined with ABC without predefined summary statistics and, for most cases, in the 50% credible intervals (dark blue). Both methods also correctly reconstructed a complex history encompassing an expansion interrupted by a bottleneck and followed by a constant size (see Figure 4 ‘Bottleneck’). However, they were unable to correctly estimate the parameters of a very complex ‘Zigzag’ history except for its initial growth period and instead reconstructed a smoother history with values intermediate to the lower and higher population sizes (see Figure 4 ’Zigzag’). This confirmed the smoothing behavior identified previously for ABC and MSMC on these demographic scenarios (Boitard et al., 2016b). Finally, similarly to ABC on predefined summary statistics (Boitard et al., 2016b), SPIDNA predictions of very recent population sizes were slightly biased toward the center of the prior distribution, however combining SPIDNA with ABC tended to correct this bias in most cases.

**Figure 4:**
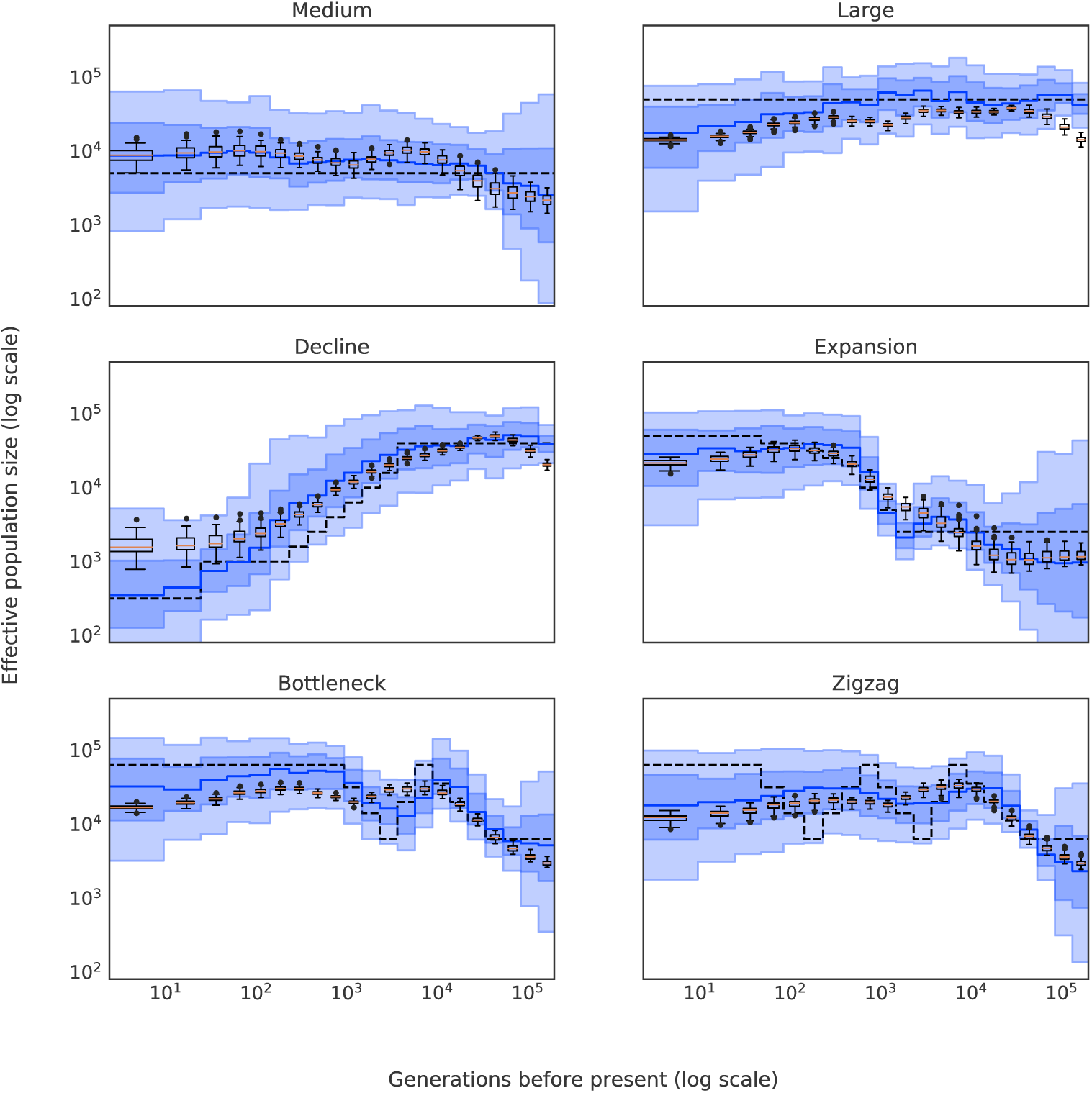
Predictions of SPIDNA and ABC using SPIDNA outputs, for six predefined scenarios (dashed black lines). 100 replicates were simulated for each scenario. Boxplots show the dispersion of SPIDNA predictions (over replicates). For each history inferred by SPIDNA combined with ABC, we display the posterior median (plain blue line), the 50% credible interval (dark blue) and the 90% credible interval (light blue).

### 3.4 Impact of positive selection on SPIDNA and ABC inference

Most inference methods are based on model assumptions that are likely to be incorrect. Violations of these assumptions have been shown to cause biases or inaccuracy in inference and simulation-based methods are similarly prone to the discrepancy between simulated space and reality: recombination, selection and heterogeneous sampling impact demographic inference (Lapierre et al., 2016, Schrider et al., 2016), population structure rather than panmixia impacts population size inference (Mazet et al., 2016), demographic uncertainty impacts selection inference (Torada et al., 2019). Here we investigated the impact of positive selection on SPIDNA and ABC inference for three illustrative demographic cases (scenarios Medium, Decline and Expansion of Figure 4). Because including selection required a change in the genetic simulator (msms instead of msprime), we first ensured that the change of tool to generate the new test dataset had no influence on the prediction accuracy (Figure S2). We then simulated 2Mb regions including a central SNP under positive selection, with varying selection strength, starting time and frequency of the beneficial allele at this time (100 regions for each scenario). We chose a conservative approach in which all 100 regions are under selection (worst case scenario). For each scenario we predicted the population size history using SPIDNA (batch normalization) or ABC (with local linear correction) on summary statistics. Both ABC and SPIDNA predictive errors varied with the selection coefficient (Figure S3). On average a moderate selective pressure (100-400) did not decrease the performance (Figure S3 top row). ABC inference for declining population datasets was the only one negatively impacted (increased error for s=200 and 400). In fact, in multiple cases increasing *s* decreased the prediction error mean. Very strong selection (*s* = 800) on the other hand led to an increased prediction error mean in all cases except for the declining histories inferred by SPIDNA. In addition, the 95% quantile and standard deviations of the prediction errors tend to increase with *s* (Figure S3) indicating that the prediction should be taken more cautiously in the presence of strong positive selection. This variance is systematically smaller for SPIDNA than ABC. In particular, a handful of histories reconstructed with ABC are far off while SPIDNA prediction errors remain comparatively low for all scenarios (Figure S4).

### 3.5 SPIDNA infers the decline of effective population size of cattle

We inferred the effective population size history of three breeds of cattle (Angus, Fleckvieh and Holstein) based on the same 75 individuals studied by Boitard et al. (2016b) and sampled by the 1,000 Bull Genomes Project (Figure 5). The best ABC and SPIDNA configurations both infer a large ancestral effective population size and a decline for the past 70,000 years. However, SPIDNA reports higher recent population sizes (Angus:11,334, Holstein:12,311, Fleckvieh:13,579) than ABC (Angus:361, Holstein:552, Fleckvieh:1,329). Interestingly, SPIDNA infers the same population sizes for all three breeds before 10 thousand years ago. This is in agreement with the estimation of the beginning of the domestication (Zeder, 2008). SPIDNA combined with ABC also reconstructed a decline after domestication but estimated larger population sizes for the last 30,000 years than SPIDNA alone. In addition Angus had the largest recent population and Fleckvieh the smallest in contrary to the two previous methods. Finally SPIDNA combined with ABC identified an episode of smooth decline and recovery of the population size preceding the domestication in the putative ancestral species (between 400,000 and 30,000 years ago). Credible intervals from Figure S6 are in accordance with the hypothesis of a decline. ABC on summary statistics did not infer this ancient change in population size (this study and Boitard et al. (2016b)), however Boitard et al. (2016a) also estimated that 123,465 years ago the ancestral population size increased from 73,042 to 137,775 using fastsimcoal2 a method based on the site-frequency-spectrum (Excoffier et al., 2013).

**Figure 5:**
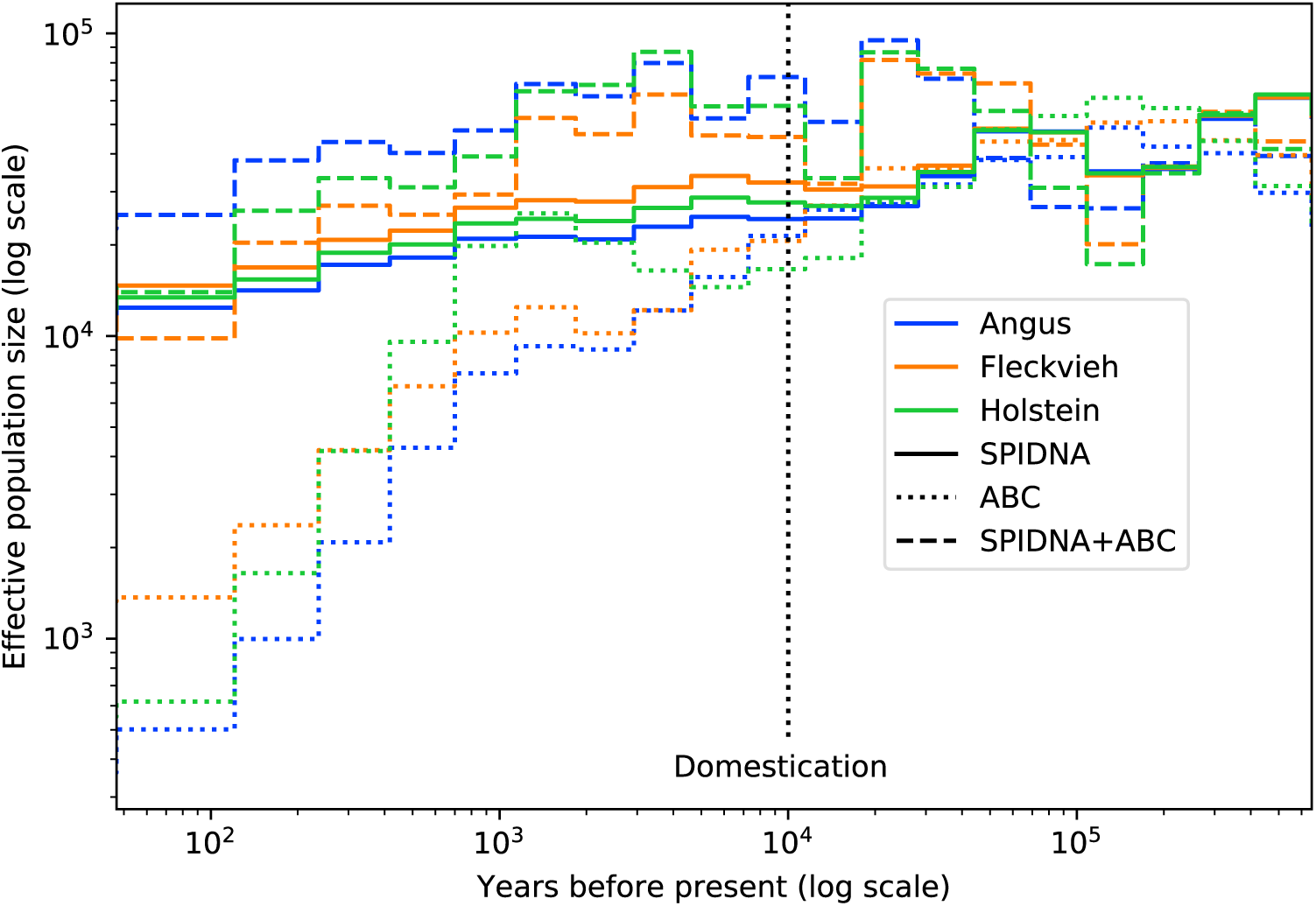
Effective population size of three cattle breeds inferred by ABC (dotted lines), by the best SPIDNA architecture, SPIDNA batch normalization (dashed lines), and by ABC based on SPIDNA outputs (dashed lines). Domestication is estimated to have occurred 10 000 years ago (vertical dotted line).

## 4 Discussion

In this paper, we introduced a deep learning approach to infer the detailed size history of a single population directly from genomic data given an unknown recombination rate. This consisted in inferring jointly 21 population size parameters. We not only increased the complexity of the demographic model with respect to previous works such as Flagel et al. (2018), but also compared the performance of our architecture to other methods including ABC, and applied our approach to real data sets. We found that our approach compared competitively with one of the best to date approaches, with the added advantage of not relying on summary statistics. A robustness analysis based on a subset of demographic scenarios also indicated that SPIDNA might be more robust than ABC to the presence of positive selection in the data. Finally, we reconstructed the effective population size fluctuations of three cattle breeds and confirmed that they all had similar sizes when they were part of the same ancestral species *Bos taurus* and underwent a decline likely linked to their domestication, although the estimated strength of this decline depended on the inference method.

### 4.1 On the practicability and importance of architecture design

When applying deep learning techniques, the design of the neural network architecture is critical, as poor design can lead to a lack of expressive power, information loss, underfitting, overfitting, or unnecessary complications that slow down the training process. The recent history of successes in Computer Vision consists in architecture improvements, leading to performance jumps (e.g. from MLP to LeNet, AlexNet, VGG, Inception and ResNet (He et al., 2016, Krizhevsky et al., 2012, LeCun et al., 1998, Simonyan and Zisserman, 2014, Szegedy et al., 2017)). But these successes have been built incrementally by relatively small changes over the last years, involving a large number of studies, researchers and challenges. Indeed finding the best architecture suited for a task is hard and time-consuming given the wide range of possibilities. Therefore, automating architecture and hyperparameter choice is an important challenge that can yield benefit to smaller fields such as population genetics. In our study, the Bayesian hyperparmeter optimisation procedure allowed us to test multiple networks thanks to a better usage of the computational power available by giving more budget to the most promising ANN architectures and hyperparameters. This procedure could be extended to hyperparameters that further describe the architecture of the network such as the number and type of layers, of neurons, the type of non-linearity or the topology. Thanks to this procedure we investigated a series of architectures, starting from the simple multi-layer fully-connected network (MLP) and moving on to more complex architectures, and exhibited the link between design and performance.

To interpret the results and compare them, let us first note that in Figure 3, a 0 error means perfect prediction, while an error of 1 means that no information is extracted from the input. Indeed, a function outputting always the same value, for all samples, can at best predict the average target value over the dataset, in which case the error is the standard deviation over the dataset of the value to predict, which is normalized to 1 in our setup.

Processing the SNP and distance matrices with a MLP led to high prediction errors, especially for recent population sizes. This is not surprising, since genomic information is encoded as a simple list of values, where the order has no meaning from the MLP point of view, which then cannot exploit information given by the data structure. In summary, an MLP configuration has several drawbacks: (i) the number of network parameters to estimate is high; (ii) the MLP can only retrieve the geometry of the data through training, with no guarantee that it will learn the spatial structure of the genome (i.e. the column order and distance between SNPs) or distinguish from which individual comes each SNP. In spite of all these hindrances, the MLP still performed far better than random guesses or constant prediction (32% better).

On the contrary, CNN layers process input elements by groups, allowing close SNPs to be processed together. This feature, combined with the stacking of layers in CNN, helps the network to construct features dependent of the SNPs proximity. Important summary statistics used in ABC or other inference methods such as linkage disequilibrium can potentially be easily expressed by such CNN. Hence we proposed several novel convolutional architectures, tailored to genetic data. We first developed a *custom* CNN with 2D filters that could have different shapes, i.e. mixed kernel sizes but also non symmetrical masks. There is indeed no rational behind considering square masks only as is usually done in computer vision to describe pixel neighborhoods, as rows and columns in our case correspond to different entities (individual or phased haplotype versus markers). Using varied mask shapes (e.g., 2×2, 5×4 or 3×8) helps our *custom* CNN to learn features of various patterns, potentially mimicking different types of summary statistics (“vertical” masks integrate over individuals, enabling the computation of allele frequencies at a SNP, while “horizontal” ones integrate over SNPs, as IBS or IBD sharing tract length does). Such mixed size filters have proved useful in the Computer Vision literature also, under the name of Inception architectures (Szegedy et al., 2017); they allow the extraction of a mixture of different kinds of information from multiple scales within the same layer. The large gap in performance between a simple MLP and this *custom* CNN confirms the importance of such considerations. A natural extension would be to integrate this feature into SPIDNA, our permutation-invariant architecture.

### 4.2 Novel architectures tailored to genomic data

#### 4.2.1 Invariance to haplotype permutation

The order in which simulated haplotypes are arranged in a SNP matrix has no meaning. Although the *custom* CNN network above cannot be guaranteed to be exactly invariant to the haplotype order, it can approximately learn this data property. To avoid wasting training time to learn that there is no information in the row order, it has been proposed to systematically sort the haplotypes according to a predefined rule (Flagel et al., 2018, Torada et al., 2019). Because there is no ordering in high dimensional space that is stable with respect to perturbations (Qi et al., 2017), we chose yet another alternative and enforced our network to be permutation-invariant by design. Permutation-invariant networks, or exchangeable networks, were successfully applied in population genetics by Chan et al. (2018) for inferring local recombination, but our architecture is different in that the invariant operations are performed at each block, enabling both individual equivariant features and global invariant features to contribute to the next layer. It has been proven that this type of architecture provides universal approximation of permutation-invariant functions (Lucas et al., 2018, Zaheer et al., 2017). Here we applied the methodology from Lucas et al. (2018) by using the mean as our invariant operation, but we encourage developers to experiment with other invariant functions such as moments of higher order. Among our permutation-invariant architectures, the best one (SPIDNA using batch normalization) had a smaller prediction error than our *custom* CNN. However, it is not clear whether this improvement is directly linked to its built-in permutation-invariance property, or to other differences between the two networks. Controlling the speed to invariance thanks to the parameter *α* improved the performance of the instance normalization SPIDNA, but not significantly the performance of the instance normalization adaptive SPIDNA (see table S2).

#### 4.2.2 Robustness to the number of individuals

Importantly, SPIDNA adapts to the number of individuals, which is an advantageous property compared to many methods relying on summary statistics. SPIDNA can be trained on data sets having similar or varying sample sizes, and, once trained, it can be directly applied to a dataset of reasonably close sample size, but unobserved during training. We provide an example of robustness experiment focusing on a few subsets of demographic scenarios (medium or large constant size populations, declining or expanding populations) and a wide range of sample sizes (from 10 to 150, Figure S5). SPIDNA using batch normalization (trained on exactly 50 individuals) did not suffer a strong loss of accuracy when the sample sizes remained in the [45,65] range. Outside of this range, the predictions were inaccurate in two cases: small sample sizes under expanding and constant size scenarios, or large sample sizes under the expansion scenario. This was expected because this specific network was not exposed to diverse sampling sizes during training. Given the observed variations across scenarios and if the sample size is expected to vary substantially from 50, we advise the user to perform a similar experiment based on her/his targeted sample size and a larger number of scenarios drawn from the prior distribution. If needed, the user can then train a new SPIDNA network without any change in its architecture, either on a set containing a wider range of sampling sizes or on a set matching the targeted sample sizes. To fasten the training, this network could be initialized with the weights of the network optimized for the sample size 50, and fine-tuned on the new set.

#### 4.2.3 Automatic adaptation to the number of SNPs

The two networks designed to be adaptive to the number of SNPs have the advantage of being applicable to genetic data of any length, the opposite of networks specific to a particular number of SNPs, which transform the data with padding or compression, or are retrained for different lengths, or take as input portions of larger sequences. Our two SPIDNA adaptive networks show results close to the best of non-adaptive versions, though slightly worse (0.469 versus 0.454, see table S2), although the difference disappears when SPIDNA is combined with ABC (0.369 versus 0.372). This small performance drop is likely due to differences in normalization rather than to the adaptive feature. Indeed, the best non-adaptive SPIDNA uses batch normalization while the adaptive versions use instance normalization as there is currently no implementation of batch normalization for batches with inputs of mixed sizes. We think that adaptive architecture could greatly benefits from an optimised implementation of adaptive batch normalization or from an implementation of batches with mixed data sizes. Nonetheless, SPIDNA networks with instance normalization had a much better performance when using all SNPs rather than the 400 first SNPs only, which suggests that adaptability is a useful feature (see table S2).

Our adaptive architecture provides an alternative to data compression based on computer vision algorithms: since compression is not optimized for the task of interest, it could induce information loss by reducing data prematurely. Note indeed that the success of deep learning in computer vision lies precisely in the replacement of ad-hoc data descriptors and processing pipelines (e.g., SIFT features to describe image keypoints (Lowe, 2004), and the “bag of visual words” pipeline (Sivic and Zisserman, 2003) to build an exploitable representation of them through clustering and histograms) by ones that can be optimized. It is also an alternative to padding, a technique that consists in completing the SNP and distance matrices at the edges so that they all match the biggest simulated SNP matrix; it is left to the neural network to guess where the real genetic data stops and where padding starts. As such it may make the task more difficult, given that the SNP matrix size is highly variable between different demographic histories and some examples would contain more padding values that actual genetic information. RNN are also a natural alternative to process sequence of variable size, though they induce an unequal contribution of SNPs to the final result, depending on their ordering along the sequence. Indeed, as the information from the previous elements of the sequence is stored in the internal state of the RNN, earlier parts of the sequence can be more easily forgotten. Nonetheless, they were very recently proven to be useful to predict local recombination rate along the genome (Adrion et al., 2019) and future works should investigate whether this scales up to global characteristics and to a different task.

### 4.3 Advantages and challenges of deep learning

Alongside the quality of deep learning to automatically extract informative features from high dimensional data, artificial neural networks are also very flexible. For instance, they can be used for transfer learning, that is, a network trained for a specific task can be reused for another one by only modifying the last layers (e.g. a network trained for population size history inference could be reused for classification between demographic scenarios) (Pan and Yang, 2009). The new network will benefit from the embedding already learned for the previous task, improving error and learning time. We also highlight that, as for most ABC methods, the parameters are inferred jointly, a major point as the common population genetics model parameters almost never have an independent impact on shaping genetic diversity. We noted that for highly fluctuating population sizes, SPIDNA estimated smooth histories. Smoothing can be seen as a good byproduct and was for example achieved on purpose by SMC++ thanks to a spline regularization scheme (Terhorst et al., 2017). A tentative explanation for SPIDNA’s smoothing effect while no regularizer was used is that it is easy for neural networks to express smoothing filters in their last layer. As, in our task, smoothing is correlated with lower prediction variance, the training of SPIDNA naturally chooses to smooth out its predictions. This could be seen as a tendency to favor low variance in the bias/variance trade-off.

#### 4.3.1 Combining deep learning and Approximate Bayesian Computation to approximate the posterior distribution

We found that adding an ABC step to the deep learning approach increased its performance. This ABC step takes as input the demographic parameters predicted by SPIDNA instead of the usual summary statistics. This strategy was proposed by Jiang et al. (2017) that showed that a deep neural network could approximate the parameter posterior means, which are desirable summary statistics for ABC. It was applied under the name of ABC-DL in two population genetics studies for performing model selection, however both papers relied on the joint SFS as predefined candidate summary statistics (Lorente-Galdos et al., 2019, Mondal et al., 2019). Here, we are taking advantage of both the deep architecture to bypass summary statistics and the Bayesian framework to refine the prediction and approximate the posterior distribution. The statistics currently processed by ABC are the average over multiple independent regions of SPIDNA predicted population sizes. A natural future step would be to investigate whether combining differently these regions lead to improved predictions.

It not yet clear why this combination decreases the prediction error. Neural networks, such as SPIDNA, learn a very general mapping of the whole input space to the output demographic parameter space. On the other hand, ABC learns a local relationship, the posterior distribution of the demographic parameters, for each targeted/observed example based on its neighbourhood in the input space. Combining ABC with SPIDNA thus adds a local inference step to the general mapping learnt by SPIDNA, and this might help readjusting the predictions locally. This is illustrated on Figure 4 where recent population sizes estimated by SPIDNA have a tendency towards the center of the prior while SPIDNA+ABC corrects it. This combination might be modifying the bias/variance trade-off favored by SPIDNA towards higher variance. These hypotheses could be investigated further in future works.

This gain however comes with a disadvantage which is the need for ABC to approximate a posterior distribution for each new dataset. This can be fairly time consuming for large panels containing many populations for which demography has to be reconstructed. Contrary to ABC, SPIDNA and other deep learning approaches, once trained, provide immediate predictions. This amortization of the training time is relevant for all studies processing large number of datasets such as meta analyses over populations or species (e.g Roux et al. (2016)) or addressing window-based tasks, such as selection and introgression scans, local ancestry or recombination estimations. In these cases the parameter predictive uncertainty could be estimated by the network (Chan et al., 2018, Lakshminarayanan et al., 2017) rather than through an ABC procedure.

Finally, we showed that applying ABC to SPIDNA predictions combined with precomputed summary statistics led to an error 4.7% smaller than the one of a regular ABC and 6.0% smaller than SPIDNA. This indicates that the information retrieved by SPIDNA does not completely overlap the one encoded into the predefined summary statistics but is not completely orthogonal either. The different behaviours of SPIDNA and ABC in term of robustness to the presence of selection also support this hypothesis. These are the first steps towards understanding and interpreting the artificial neural networks currently used in population genetics, a major challenge that the deep learning field currently faces for many of its applications (Gilpin et al., 2018) and that has not yet been investigated in our community.

#### 4.3.2 Application to real data

Applying a method trained on simulated data to a real dataset can be a difficult task. Here we show that the estimated effective population sizes of the three cattle breeds were qualitatively similar across the different methods used. All of them were able to recover the large ancestral population size shared by the three breeds, followed by its decline after domestication. However, the methods produced size estimates that were quantitatively different, notably in the strength of the decline and the recent population sizes. For quality reasons, inference was done using genotypes pruned of low frequency alleles rather than haplotypes. The architecture and hyperparameters were optimized based on simulated haplotypes, and the network was trained on simulated genotypes. It is possible that an architecture designed with a new hyperparameter optimization procedure calibrated for filtered genotypes would decrease SPIDNA error rate. However, the discrepancy between ABC and SPIDNA reconstructions in the last 10,000 years might also be due to the sensitivity of ANNs to overfitting and to mispecifications in the model generating training data. For example, decrease in performance due to demographic mispecification has already been shown for selection inference based on ANNs (Torada et al., 2019). In our work we investigated whether positive selection on de-novo mutation or standing variation could have such a strong effect on demographic inference and found that SPIDNA was robust to various selective scenarios. In the cattle case, model mispecification arises because cattle breeds are subjected to strong artificial selection pressures based on observed phenotypes, with few males contributing to the next generations, which is an extreme case of selection and a clear violation of the coalescent assumptions underlying our training simulated set. In addition, errors or missing information in real data were not modelled in the training set, a procedure that can improve ABC performance when using multiple summary statistics such as haplotype length statistics (Jay et al., 2019). When comparing performance on training and validation sets, we found that our architectures were not overfitting. Yet it is possible that the features automatically constructed by ANNs are more sensitive to a gap between real and simulated data (e.g. unmodelled errors and artificial selection) than an ABC method based on SFS and LD statistics. Although we checked the robustness of SPIDNA to the simulator tool and to multiple cases of positive selection on haplotype data (Figures S2 and S3), artificial selection based on phenotype and pedigree information is yet another type of model violation. Systematically testing and improving the robustness of ANNs trained on simulations is a great challenge for the coming years.

## Conclusion

We addressed a challenging task in population genetics, that is, reconstructing effective population size through time. We showed that this demographic inference could be done for unknown recombination rates. Our approach has only a slight increase in performance compared to the more classical method (ABC based on summary statistics) albeit it does not require any expert knowledge regarding the computation of summary statistics. Besides, the ABC approach (without predefined statistics) can be combined to obtain posterior distributions. We are confident that a network exchangeable and adaptive to the input size is a promising architecture for future lines of works for other population genetics tasks, as it could prevent premature loss of information and favor learning new features rather than known haplotype invariance. These new features can be seen as automatically learned summary statistics and will be crucial in areas where inference is challenging and for which researchers are always designing novel and hopefully expressive summary statistics (see for example the recent line of research on adaptive introgression (Racimo et al., 2015)). As for now, co-estimating multiple processes remains a hard task, and inference is mostly done under a simplifying assumption, e.g. selection or recombination are inferred under a fixed demographic scenario and step-wise population size is reconstructed for a single panmictic population (but see MSMC and MSMC-IM for an extension to two populations with interactions (Schiffels and Durbin, 2014, Wang et al., 2019)). The success of ABC and simulation-based methods is partly due to their convenience to include complex models via simulations. Here we showed, for the first time, that a well designed artificial neural network is capable of retrieving information about fluctuating effective population size, competes favorably with a commonly used approach, and can also be combined with existing summary statistics if needed. Additionally, recent studies showed that artificial neural networks could detect introgression and selection (Flagel et al., 2018, Torada et al., 2019). For the above reasons, and because extracting information automatically should lead to the identification of features that disentangle processes hardly distinguishable, we are hopeful that future robust networks trained on complex simulations could help solving jointly some of these tasks. Finally, we provided (i) a tool for users willing to infer population size history of any species that can be applied to phased or unphased genomes (available from https://gitlab.inria.fr/ml_genetics/public/dlpopsize); (ii) new exchangeable network architectures, some of which have the promising feature of being adaptive to input size; (iii) guidelines for future developers on building architectures and hyper-optimization to facilitate the development of new artificial neural networks for population genomics.

## Acknowledgments

We are grateful to the genotoul bioinformatics platform Toulouse Midi-Pyrenees (Bioinfo Genotoul) and the TAU team for providing computing and storage resources, to the Paris-Saclay Center for Data Science 2.0 (IRS) for funding. JC salary was funded by DIM-1Health. We are also grateful to Simon Boitard for helpful discussions and providing the cattle dataset. We thank Michael Blum for his comments on a first version of the manuscript and Diviyan Kalainathan for his support with the Titanic platform. We also thank Jeffrey Spence, Lex Flagel and two anonymous reviewers for their comments.

## Data Availability Statement

The data that support the findings of this study are openly available in *dlpopsize* at https://gitlab.inria.fr/ml_genetics/public/dlpopsize.

## Supplementary Materials

### Computational resources

Simulations have been performed on the genotoul bioinformatics platform with the following hardware:

- 68 nodes with 2 E5-2670 v2 Intel CPUs (2.50GHz, 20 threads) and 256GB of RAM
- 48 nodes with 2 E5-2683 v4 Intel CPUs (2.10GHz, 32 threads) and 256GB of RAM.

All summary statistics, trainings and predictions were computed on the TAU’s Titanic platform with the following hardware:

- 5 nodes with 4 GTX 1080 (12GB of VRAM) GPUs, 2 E5-2650 v4 Intel CPUs (2.20GHz, 24 threads) and 252GB of RAM
- 7 nodes with 4 RTX 2080 (12GB of VRAM) GPUs, 2 Silver 4108 Intel CPUs (1.80GHz, 18 threads) and 252GB of RAM
- 1 node with 4 Tesla P100 (16GB of VRAM) GPUs, 2 E5-2690 v4 Intel CPUs (2.60GHz, 28 threads) and 252GB of RAM
- 1 node with 2 RTX 2080 (8GB of VRAM) GPUs, 2 E5-2650 v4 Intel CPUs (2.20GHz, 24 threads) and 252GB of RAM

Both platforms use Slurm as job scheduling system. Batch sizes and deep learning architectures were all designed to fit on less than 12GB of VRAM during training. To train non-adaptive architectures, batches were split between 3 GPUs with at least 12GB of VRAM. Adaptive architectures were trained on one GPUs as batch data of varying sizes could not be concatenated in the same tensor. The training of SPIDNA took at most 1h42 per epoch for non-adaptive version and 31h31 per epoch for adaptive version. The slow computation time of adaptive SPIDNA is mostly due to data being inputted one by one in the network instead of concatenated in tensors.

## Supplementary figures and tables

**Figure S1:**
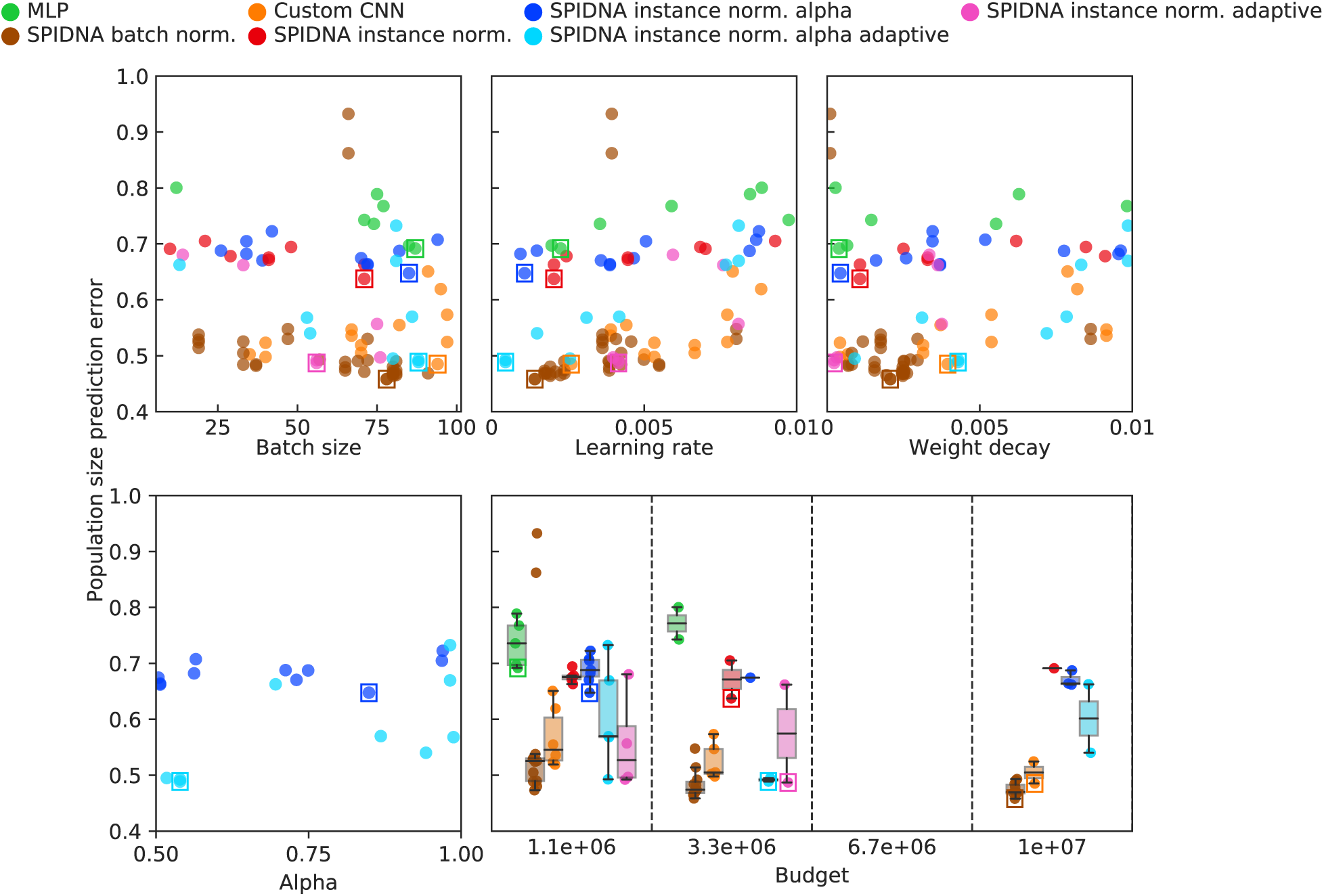
Population size prediction error for each run of the hyperparameter optimization procedure. X-axes indicate the hyperparameter (batch size, learning rate, weight decay and alpha) or budget values, and colors indicate the type of network used for the run (MLP, *custom* CNN and multiple SPIDNA architectures). For each network the best run is surrounded by a square.

**Figure S2:**
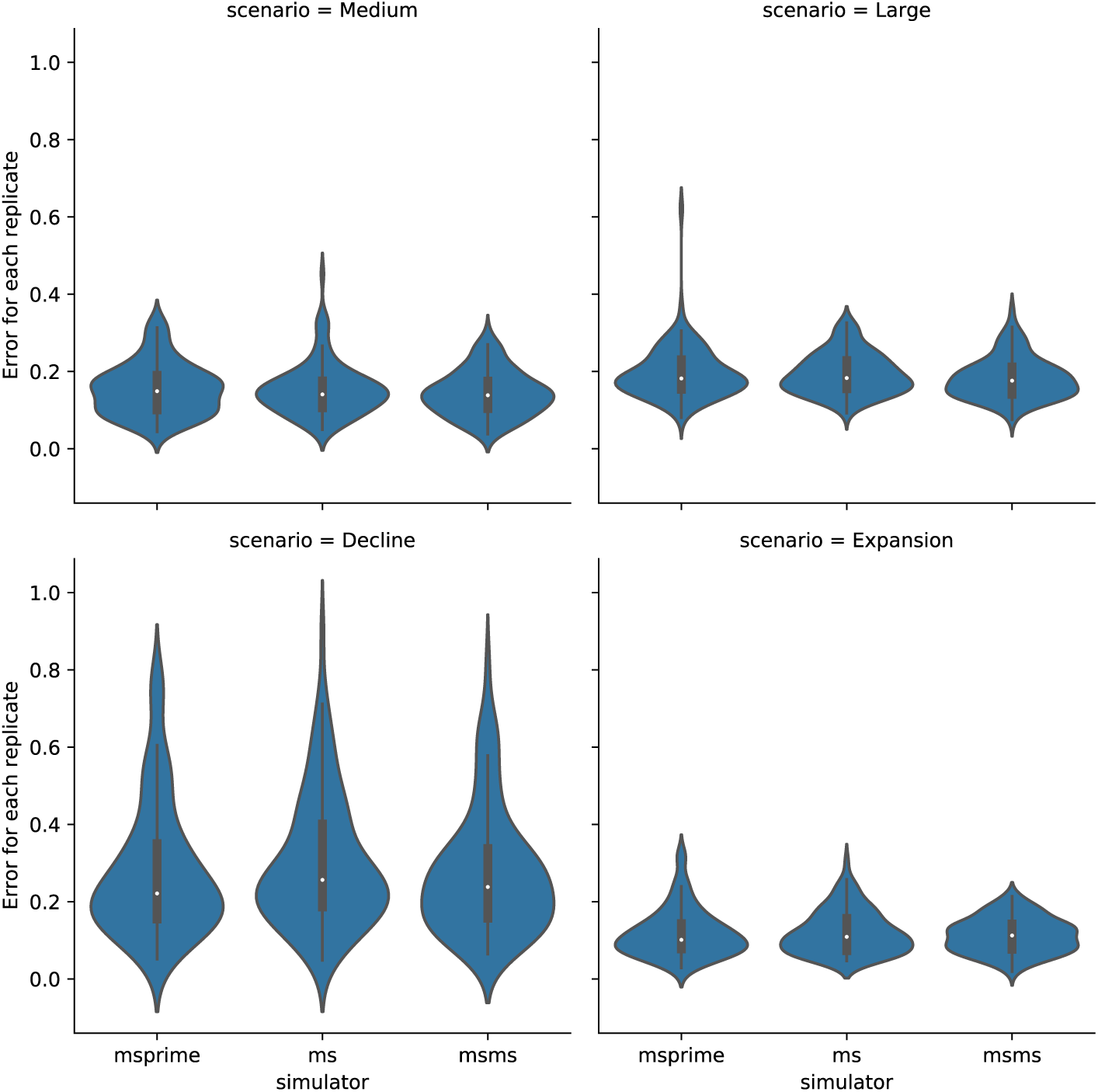
Robustness to simulator tool. Distributions of SPIDNA predictive errors per replicate (i.e, per independent genomic region) for four demographic scenarios and three different genetic simulators (msprime, ms, msms). SPIDNA batch norm. network was trained on simulated datasets generated with msprime. The test datasets were generated by different simulators, based on the same demographic parameters and under neutrality. X-axes: simulator for the test set; Y-axes: predictive error for each region/replicate (i.e. for each matrix of size 50 samples × 400 SNPs) averaged over the 21 time steps. Each violin describes 100 replicates.

**Figure S3:**
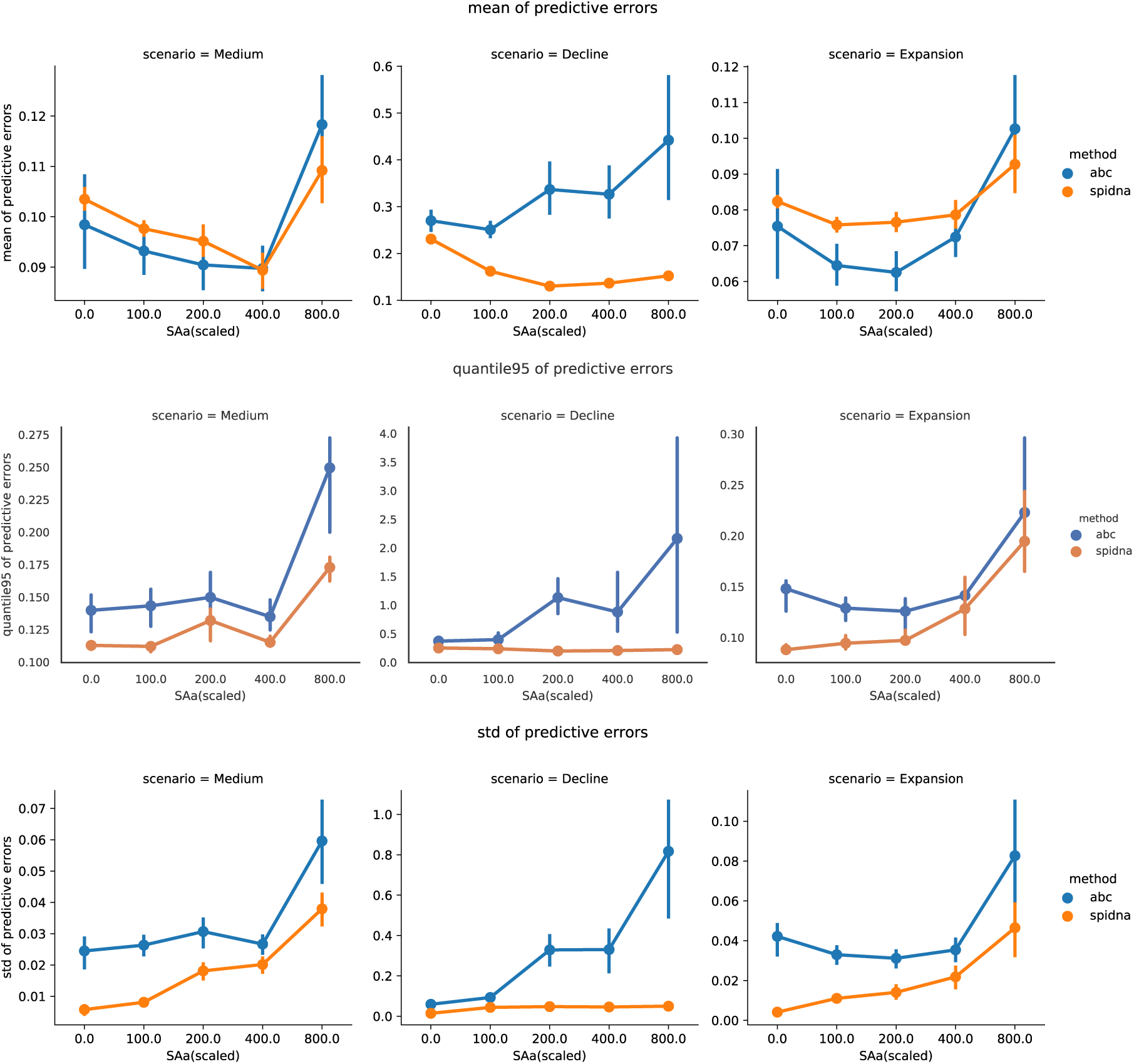
Robustness to the presence of positive selection. ABC and SPIDNA (batch norm.) predictive errors computed from 100 2Mb-long regions for three demographic scenarios (Medium constant size, Decline and Expansion) under various selective pressures (with additive fitness effect). The reference/training set is the same as the one used throughout the paper (neutral simulations generated with msprime from a prior distribution on recombination rate and population sizes). The test datasets were simulated using msms with multiple values of selection strength, starting time and initial frequency of the beneficial allele. X-axes: Selection coefficient SAa. Y-axes: Mean (top), 95% quantile (middle row) and variance (bottom row) estimators of the predictive error (across 30 test sets for SAa=0 and 144 test sets for any other SAa value). Vertical bars correspond to 95% confidence intervals computed via bootstrap.

**Figure S4:**
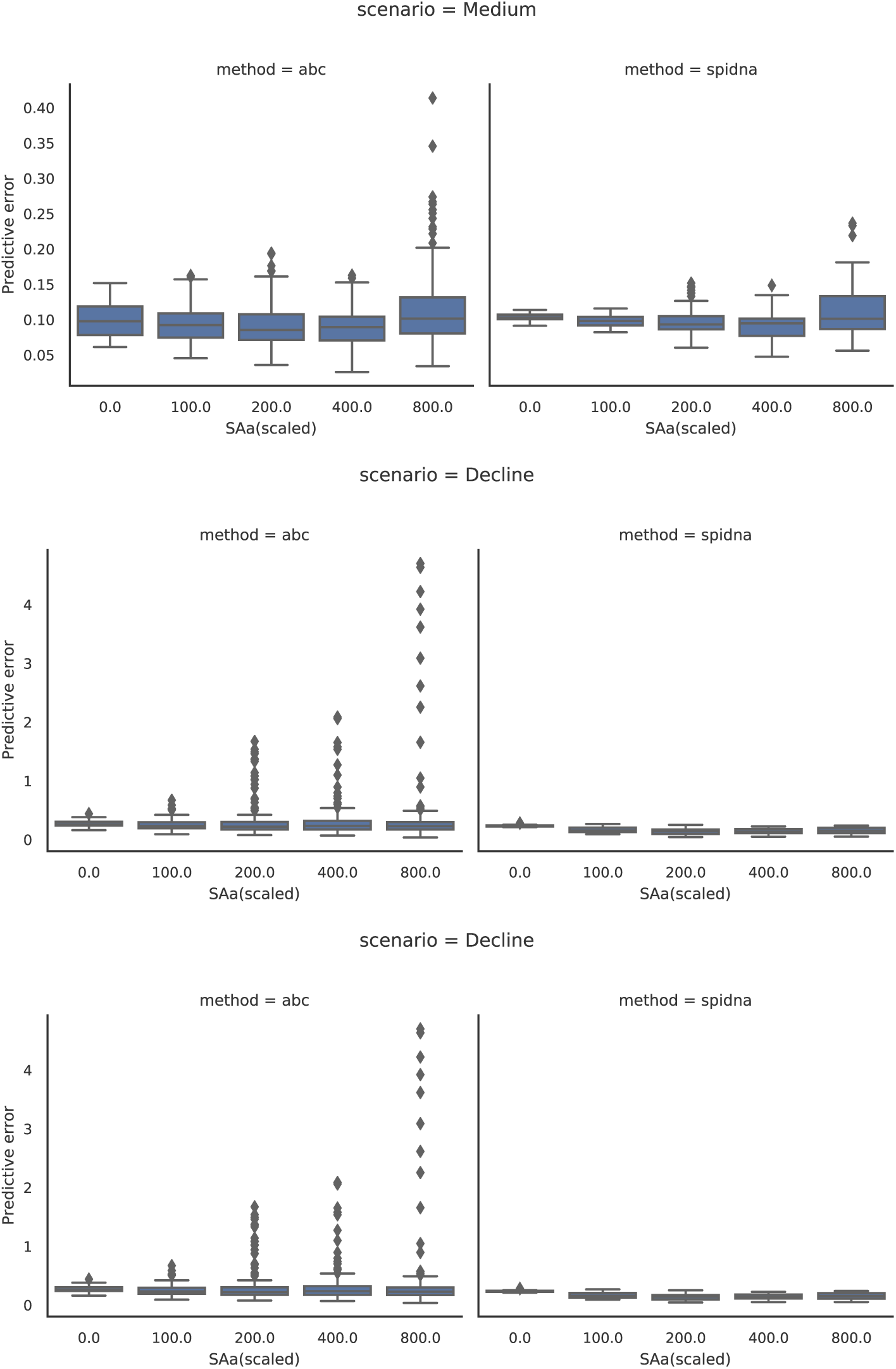
Robustness to the presence of positive selection. ABC and SPIDNA (batch norm.) predictive errors computed from 100 2Mb-long regions for three demographic scenarios (Medium constant size, Decline and Expansion) under various selective pressures. The reference/training set is the same as the one used throughout the paper (neutral simulations generated with msprime from a prior distribution on recombination rate and population sizes). The test datasets were simulated using msms with multiple values of selection strength, starting time and initial frequency of the beneficial allele. X-axes: Selection coefficient SAa. Y-axes: Distribution of predictive errors (across 30 test sets for *SAa* = 0 and 144 test sets for any other SAa value).

**Figure S5:**
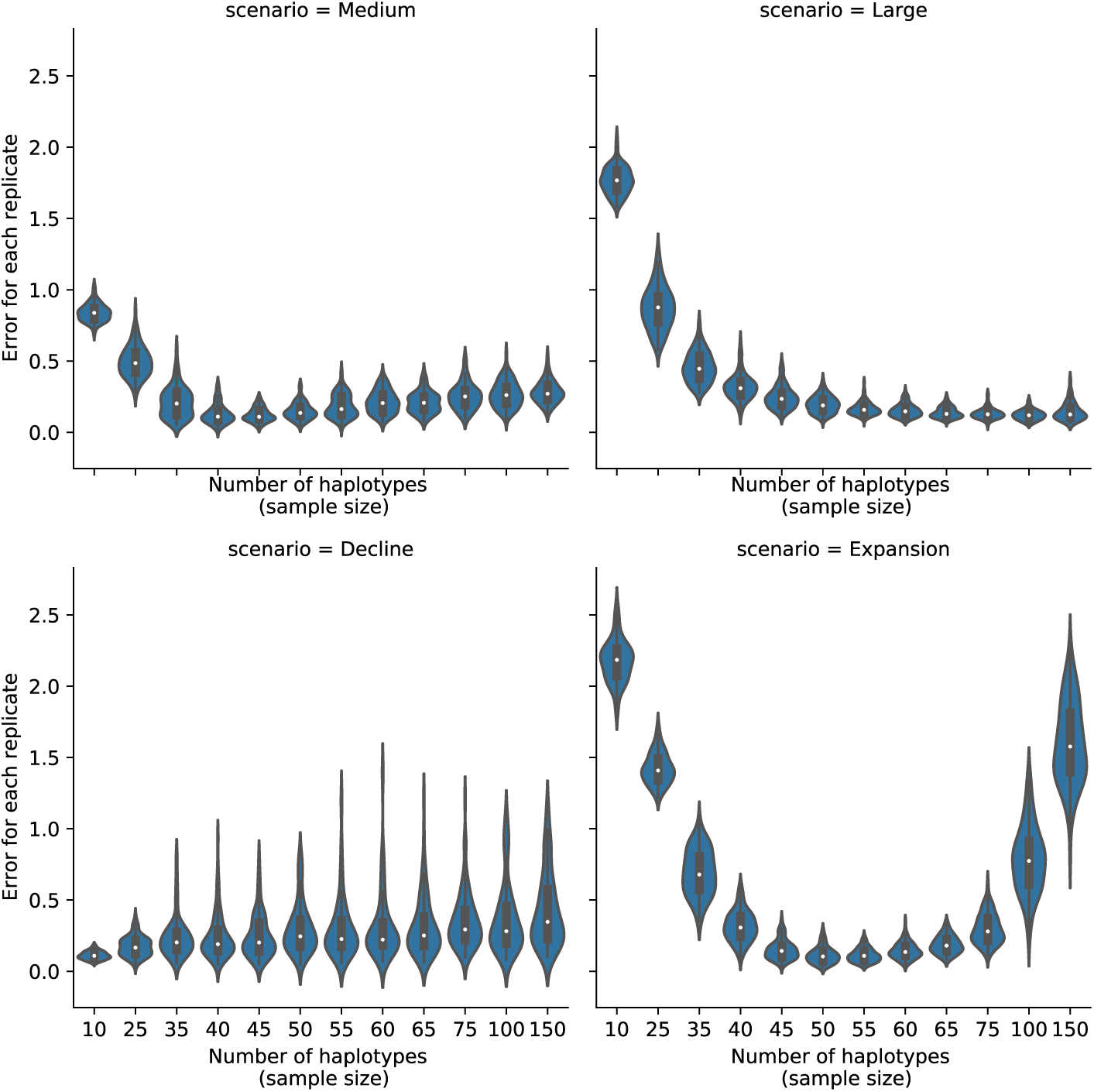
Robustness to sample size. Distributions of SPIDNA predictive errors per replicate (i.e, per independent genomic region) for four demographic scenarios and different sampling sizes. SPIDNA (batch norm.) network was trained on simulated datasets containing exactly 50 samples. The test datasets were simulated with msprime based on the same four demographic parameter sets but with different sample sizes (ranging from 10 to 150 haplotypes). X-axes: sample size *M* of the targeted region; Y-axes: predictive error for each replicate (i.e. for each matrix of size *M* samples ×400 SNPs) averaged over the 21 time steps. Each violin describes 100 replicates.

**Figure S6:**
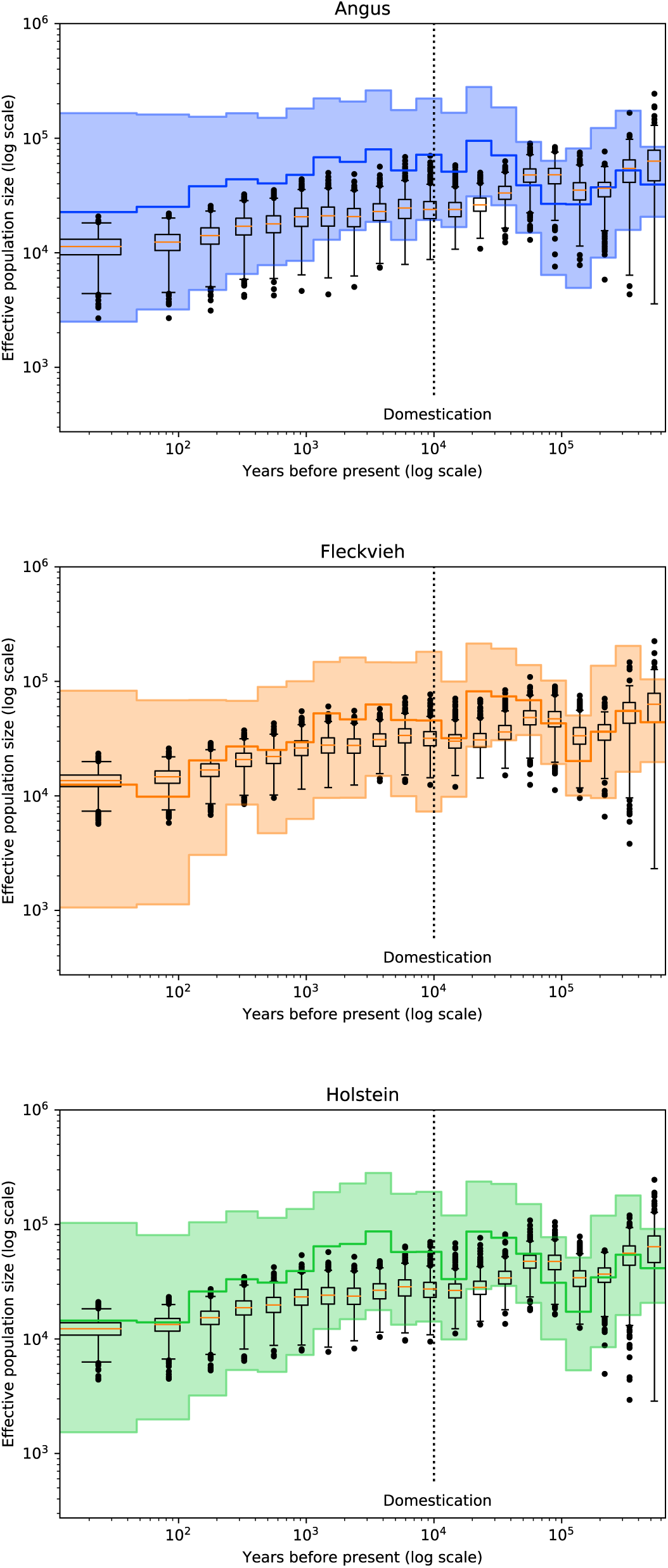
Effective population size of three cattle breeds inferred by the best SPIDNA architecture, SPIDNA batch normalization, and by ABC based on SPIDNA outputs. Boxplots show the dispersion of SPIDNA predictions (over replicates). For each history inferred by SPIDNA combined with ABC, we display the posterior median (plain line) and the 95% credible interval. Domestication is estimated to have occurred 10 000 years ago (vertical dotted line).

**Table S1:** Parameters used for the *Flagel* CNN

| input dimension | SNP encoding | Convolution type | Kernel size | Pooling size | Log-scaled output? | Sort chromosomes? | Use dropout? |
| --- | --- | --- | --- | --- | --- | --- | --- |
| $50 \times 400$ | 0/-1 | 1D | 2 | 2 | Yes | Yes | Yes |
| $50 \times 1784$ | 0/-1 | 1D | 2 | 2 | Yes | Yes | Yes |
| $50 \times 400$ | -1/1 | 1D | 2 | 2 | Yes | Yes | No |
| $50 \times 1784$ | -1/1 | 1D | 2 | 2 | Yes | Yes | No |

**Table S2:** Prediction errors of the best configuration of each method after hyperparameter optimization

|  | Method | Adaptive | Summary statistics | ABC correction | Alpha | Validation error | Test error |
| --- | --- | --- | --- | --- | --- | --- | --- |
| 0 | ABC | No | Yes | No | No | 0.490 | 0.496 |
| 1 | ABC | No | Yes | Linear reg. | No | 0.357 | 0.369 |
| 2 | ABC | No | Yes | Ridge reg. | No | 0.363 | 0.376 |
| 3 | ABC | No | Yes | Single layer NN | No | 0.352 | 0.364 |
| 4 | MLP | No | Yes | No | No | 0.399 | 0.437 |
| 5 | MLP | No | No | No | No | 0.690 | 0.675 |
| 6 | Custom CNN | No | No | No | No | 0.485 | 0.487 |
| 7 | Flagel CNN 0/-1 encoding | No | No | No | No | 0.537 | 0.541 |
| 8 | Flagel CNN 0/-1 encoding | Downsampling | No | No | No | 0.437 | 0.444 |
| 9 | Flagel CNN -1/1 encoding | No | No | No | No | 0.610 | 0.609 |
| 10 | Flagel CNN -1/1 encoding | Downsampling | No | No | No | 0.482 | 0.484 |
| 11 | SPIDNA | No | No | No | No | 0.453 | 0.454 |
| 12 | SPIDNA | No | No | No | No | 0.637 | 0.641 |
| 13 | SPIDNA | Yes | No | No | No | 0.487 | 0.489 |
| 14 | SPIDNA | No | No | No | 0.849 | 0.592 | 0.599 |
| 15 | SPIDNA | Yes | No | No | 0.539 | 0.466 | 0.469 |
| 16 | ABC + SPIDNA | No | No | No | No | 0.462 | 0.462 |
| 17 | ABC + SPIDNA | No | No | Linear reg. | No | 0.364 | 0.377 |
| 18 | ABC + SPIDNA | No | No | Ridge reg. | No | 0.371 | 0.380 |
| 19 | ABC + SPIDNA | No | No | Single layer NN | No | 0.363 | 0.372 |
| 20 | ABC + SPIDNA | Yes | No | No | 0.539 | 0.458 | 0.460 |
| 21 | ABC + SPIDNA | Yes | No | Linear reg. | 0.539 | 0.363 | 0.369 |
| 22 | ABC + SPIDNA | Yes | No | Ridge reg. | 0.539 | 0.382 | 0.391 |
| 23 | ABC + SPIDNA | Yes | No | Single layer NN | 0.539 | 0.374 | 0.384 |
| 24 | ABC + SPIDNA | No | Yes | No | No | 0.476 | 0.478 |
| 25 | ABC + SPIDNA | No | Yes | Linear reg. | No | 0.339 | 0.353 |
| 26 | ABC + SPIDNA | No | Yes | Ridge reg. | No | 0.341 | 0.357 |
| 27 | ABC + SPIDNA | No | Yes | Single layer NN | No | 0.345 | 0.361 |
| 28 | ABC + SPIDNA | Yes | Yes | No | 0.539 | 0.474 | 0.478 |
| **29** | **ABC + SPIDNA** | **Yes** | **Yes** | **Linear reg.** | **0.539** | **0.335** | **0.347** |
| 30 | ABC + SPIDNA | Yes | Yes | Ridge reg. | 0.539 | 0.339 | 0.354 |
| 31 | ABC + SPIDNA | Yes | Yes | Single layer NN | 0.539 | 0.347 | 0.365 |

## References

Jeffrey R Adrion, Jared G Galloway, and Andrew D Kern. Inferring the landscape of recombination using recurrent neural networks. bioRxiv, page 662247, 2019.

Simon Aeschbacher, Mark A Beaumont, and Andreas Futschik. A novel approach for choosing summary statistics in approximate bayesian computation. Genetics, 192(3):1027–1047, 2012.

Babak Alipanahi, Andrew Delong, Matthew T Weirauch, and Brendan J Frey. Predicting the sequence specificities of dna-and rna-binding proteins by deep learning. Nature biotechnology, 33(8):831, 2015.

Y. Bengio, P. Simard, and P. Frasconi. Learning long-term dependencies with gradient descent is difficult. Trans. Neur. Netw., 5(2):157–166, March 1994. ISSN 1045-9227. doi: 10.1109/72.279181. URL https://doi.org/10.1109/72.279181.

Anders Bergström, Shane A McCarthy, Ruoyun Hui, Mohamed A Almarri, Qasim Ayub, Petr Danecek, Yuan Chen, Sabine Felkel, Pille Hallast, Jack Kamm, et al. Insights into human genetic variation and population history from 929 diverse genomes. bioRxiv, page 674986, 2019.

Anand Bhaskar, YX Rachel Wang, and Yun S Song. Efficient inference of population size histories and locus-specific mutation rates from large-sample genomic variation data. Genome research, 25(2):268–279, 2015.

Michael GB Blum. Approximate bayesian computation: a nonparametric perspective. Journal of the American Statistical Association, 105(491):1178–1187, 2010.

Michael GB Blum, Maria Antonieta Nunes, Dennis Prangle, Scott A Sisson, et al. A comparative review of dimension reduction methods in approximate bayesian computation. Statistical Science, 28(2):189–208, 2013.

Simon Boitard, Mekki Boussaha, Aurélien Capitan, Dominique Rocha, and Bertrand Servin. Uncovering adaptation from sequence data: lessons from genome resequencing of four cattle breeds. Genetics, 203(1):433–450, 2016a.

Simon Boitard, Willy Rodriguez, Flora Jay, Stefano Mona, and Frédéric Austerlitz. Inferring population size history from large samples of genome-wide molecular data-an approximate bayesian computation approach. PLoS genetics, 12(3):e1005877, 2016b.

Michael Bridges, Elizabeth A Heron, Colm O’Dushlaine, Ricardo Segurado, Derek Morris, Aiden Corvin, Michael Gill, Carlos Pinto, International Schizophrenia Consortium, et al. Genetic classification of populations using supervised learning. PloS one, 6(5), 2011.

Jeffrey Chan, Valerio Perrone, Jeffrey Spence, Paul Jenkins, Sara Mathieson, and Yun Song. A likelihood-free inference framework for population genetic data using exchangeable neural networks. In Advances in Neural Information Processing Systems, pages 8594–8605, 2018.

1000 Genomes Project Consortium et al. A map of human genome variation from population-scale sequencing. Nature, 467(7319):1061, 2010.

1000 Genomes Project Consortium et al. A global reference for human genetic variation. Nature, 526(7571):68, 2015.

Katalin Csilléry, Olivier François, and Michael GB Blum. abc: an r package for approximate bayesian computation (abc). Methods in ecology and evolution, 3(3):475–479, 2012.

Hans D Daetwyler, Aurélien Capitan, Hubert Pausch, Paul Stothard, Rianne Van Binsbergen, Rasmus F Brøndum, Xiaoping Liao, Anis Djari, Sabrina C Rodriguez, Cécile Grohs, et al. Whole-genome sequencing of 234 bulls facilitates mapping of monogenic and complex traits in cattle. Nature genetics, 46(8):858, 2014.

Gregory Ewing and Joachim Hermisson. Msms: a coalescent simulation program including recombination, demographic structure and selection at a single locus. Bioinformatics, 26(16):2064–2065, 2010.

Laurent Excoffier, Isabelle Dupanloup, Emilia Huerta-Sánchez, Vitor C Sousa, and Matthieu Foll. Robust demographic inference from genomic and snp data. PLoS genetics, 9(10):e1003905, 2013.

Stefan Falkner, Aaron Klein, and Frank Hutter. BOHB: Robust and efficient hyperparameter optimization at scale. In Jennifer Dy and Andreas Krause, editors, Proceedings of the 35th International Conference on Machine Learning, volume 80 of Proceedings of Machine Learning Research, pages 1437-1446, Stockholmsmässan, Stockholm Sweden, 10–15 Jul 2018. PMLR. URL http://proceedings.mlr.press/v80/falkner18a.html.

Paul Fearnhead and Dennis Prangle. Constructing summary statistics for approximate bayesian computation: semiautomatic approximate bayesian computation. Journal of the Royal Statistical Society: Series B (Statistical Methodology), 74(3):419–474, 2012.

Lex Flagel, Yaniv Brandvain, and Daniel R Schrider. The unreasonable effectiveness of convolutional neural networks in population genetic inference. Molecular biology and evolution, 36(2):220–238, 2018.

Leilani H Gilpin, David Bau, Ben Z Yuan, Ayesha Bajwa, Michael Specter, and Lalana Kagal. Explaining explanations: An overview of interpretability of machine learning. In 2018 IEEE 5th International Conference on data science and advanced analytics (DSAA), pages 80–89. IEEE, 2018.

Ariella L Gladstein and Michael F Hammer. Substructured population growth in the ashkenazi jews inferred with approximate bayesian computation. Molecular biology and evolution, 36(6):1162–1171, 2019.

Xavier Glorot and Yoshua Bengio. Understanding the difficulty of training deep feedforward neural networks. In In Proceedings of the International Conference on Artificial Intelligence and Statistics (AISTATS’10). Society for Artificial Intelligence and Statistics, 2010.

Kaiming He, Xiangyu Zhang, Shaoqing Ren, and Jian Sun. Deep residual learning for image recognition. In Proceedings of the IEEE conference on computer vision and pattern recognition, pages 770–778, 2016.

Simon YW Ho and Beth Shapiro. Skyline-plot methods for estimating demographic history from nucleotide sequences. Molecular ecology resources, 11(3):423–434, 2011.

Kishore Jaganathan, Sofia Kyriazopoulou Panagiotopoulou, Jeremy F McRae, Siavash Fazel Darbandi, David Knowles, Yang I Li, Jack A Kosmicki, Juan Arbelaez, Wenwu Cui, Grace B Schwartz, et al. Predicting splicing from primary sequence with deep learning. Cell, 176(3):535–548, 2019.

Flora Jay, Simon Boitard, and Frédéric Austerlitz. An abc method for whole-genome sequence data: inferring paleolithic and neolithic human expansions. Molecular biology and evolution, 36(7):1565–1579, 2019.

Bai Jiang, Tung-yu Wu, Charles Zheng, and Wing H Wong. Learning summary statistic for approximate bayesian computation via deep neural network. Statistica Sinica, pages 1595–1618, 2017.

Jerome Kelleher, Alison M Etheridge, and Gilean McVean. Efficient coalescent simulation and genealogical analysis for large sample sizes. PLoS Comput Biol, 12(5):1–22, 05 2016. doi: 10.1371/journal.pcbi.1004842. URL http://dx.doi.org/10.1371%2Fjournal.pcbi.1004842.

Diederik P. Kingma and Jimmy Ba. Adam: A method for stochastic optimization, 2014.

Alex Krizhevsky, Ilya Sutskever, and Geoffrey E Hinton. Imagenet classification with deep convolutional neural networks. In Advances in neural information processing systems, pages 1097–1105, 2012.

Balaji Lakshminarayanan, Alexander Pritzel, and Charles Blundell. Simple and scalable predictive uncertainty estimation using deep ensembles. In Advances in neural information processing systems, pages 6402–6413, 2017.

Marguerite Lapierre, Camille Blin, Amaury Lambert, Guillaume Achaz, and Eduardo PC Rocha. The impact of selection, gene conversion, and biased sampling on the assessment of microbial demography. Molecular biology and evolution, 33(7):1711–1725, 2016.

Yann LeCun, Yoshua Bengio, et al. Convolutional networks for images, speech, and time series. The handbook of brain theory and neural networks, 3361(10):1995, 1995.

Yann LeCun, Léon Bottou, Yoshua Bengio, Patrick Haffner, et al. Gradient-based learning applied to document recognition. Proceedings of the IEEE, 86(11):2278–2324, 1998.

Liis Leitsalu, Toomas Haller, Tõnu Esko, Mari-Liis Tammesoo, Helene Alavere, Harold Snieder, Markus Perola, Pauline C Ng, Reedik Mägi, Lili Milani, et al. Cohort profile: Estonian biobank of the estonian genome center, university of tartu. International journal of epidemiology, 44(4):1137–1147, 2014.

Heng Li and Richard Durbin. Inference of human population history from individual whole-genome sequences. Nature, 475(7357):493, 2011.

Lisha Li, Kevin Jamieson, Giulia DeSalvo, Afshin Rostamizadeh, and Ameet Talwalkar. Hyperband: A novel bandit-based approach to hyperparameter optimization. arXiv preprint arXiv:1603.06560, 2016.

Xiaoming Liu and Yun-Xin Fu. Exploring population size changes using snp frequency spectra. Nature genetics, 47(5): 555, 2015.

Belen Lorente-Galdos, Oscar Lao, Gerard Serra-Vidal, Gabriel Santpere, Lukas FK Kuderna, Lara R Arauna, Karima Fadhlaoui-Zid, Ville N Pimenoff, Himla Soodyall, Pierre Zalloua, et al. Whole-genome sequence analysis of a pan african set of samples reveals archaic gene flow from an extinct basal population of modern humans into sub-saharan populations. Genome biology, 20(1):77, 2019.

David G Lowe. Distinctive image features from scale-invariant keypoints. International journal of computer vision, 60 (2):91–110, 2004.

Thomas Lucas, Corentin Tallec, Yann Ollivier, and Jakob Verbeek. Mixed batches and symmetric discriminators for GAN training. In Jennifer Dy and Andreas Krause, editors, Proceedings of the 35th International Conference on Machine Learning, volume 80 of Proceedings of Machine Learning Research, pages 2844-2853, Stockholmsmässan, Stockholm Sweden, 10–15 Jul 2018. PMLR. URL http://proceedings.mlr.press/v80/lucas18a.html.

Wenlong Ma, Zhixu Qiu, Jie Song, Jiajia Li, Qian Cheng, Jingjing Zhai, and Chuang Ma. A deep convolutional neural network approach for predicting phenotypes from genotypes. Planta, 248(5):1307–1318, 2018.

Iona M. MacLeod, Denis M. Larkin, Harris A. Lewin, Ben J. Hayes, and Mike E. Goddard. Inferring Demography from Runs of Homozygosity in Whole-Genome Sequence, with Correction for Sequence Errors. Molecular Biology and Evolution, 30(9):2209–2223, 07 2013. ISSN 0737-4038. doi: 10.1093/molbev/mst125. URL https://doi.org/10.1093/molbev/mst125.

Swapan Mallick, Heng Li, Mark Lipson, Iain Mathieson, Melissa Gymrek, Fernando Racimo, Mengyao Zhao, Niru Chennagiri, Susanne Nordenfelt, Arti Tandon, et al. The simons genome diversity project: 300 genomes from 142 diverse populations. Nature, 538(7624):201, 2016.

Olivier Mazet, Willy Rodríguez, Simona Grusea, Simon Boitard, and Lounès Chikhi. On the importance of being structured: instantaneous coalescence rates and human evolution—lessons for ancestral population size inference? Heredity, 116(4):362, 2016.

Alistair Miles, Peter Ralph, Summer Rae, and Rahul Pisupati. cggh/scikit-allel: v1.2.1, June 2019. URL https://doi.org/10.5281/zenodo.3238280.

Mayukh Mondal, Jaume Bertranpetit, and Oscar Lao. Approximate bayesian computation with deep learning supports a third archaic introgression in asia and oceania. Nature communications, 10(1):246, 2019.

Shigeki Nakagome, Kenji Fukumizu, and Shuhei Mano. Kernel approximate bayesian computation in population genetic inferences. Statistical applications in genetics and molecular biology, 12(6):667–678, 2013.

Miguel Navascués, Raphaël Leblois, and Concetta Burgarella. Demographic inference through approximate-bayesian-computation skyline plots. PeerJ, 5:e3530, 2017.

Luca Pagani, Daniel John Lawson, Evelyn Jagoda, Alexander Mörseburg, Anders Eriksson, Mario Mitt, Florian Clemente, Georgi Hudjashov, Michael DeGiorgio, Lauri Saag, et al. Genomic analyses inform on migration events during the peopling of eurasia. Nature, 538(7624):238, 2016.

Sinno Jialin Pan and Qiang Yang. A survey on transfer learning. IEEE Transactions on knowledge and data engineering, 22(10):1345–1359, 2009.

Javier Prado-Martinez, Peter H Sudmant, Jeffrey M Kidd, Heng Li, Joanna L Kelley, Belen Lorente-Galdos, Krishna R Veeramah, August E Woerner, Timothy D O’Connor, Gabriel Santpere, et al. Great ape genetic diversity and population history. Nature, 499(7459):471, 2013.

Charles R Qi, Hao Su, Kaichun Mo, and Leonidas J Guibas. Pointnet: Deep learning on point sets for 3d classification and segmentation. In Proceedings of the IEEE Conference on Computer Vision and Pattern Recognition, pages 652–660, 2017.

Fernando Racimo, Sriram Sankararaman, Rasmus Nielsen, and Emilia Huerta-Sánchez. Evidence for archaic adaptive introgression in humans. Nature Reviews Genetics, 16(6):359–371, 2015.

Louis Raynal, Jean-Michel Marin, Pierre Pudlo, Mathieu Ribatet, Christian P Robert, and Arnaud Estoup. Abc random forests for bayesian parameter inference. Bioinformatics, 35(10):1720–1728, 2018.

Camille Roux, Christelle Fraisse, Jonathan Romiguier, Yoann Anciaux, Nicolas Galtier, and Nicolas Bierne. Shedding light on the grey zone of speciation along a continuum of genomic divergence. PLoS biology, 14(12), 2016.

David E. Rumelhart, Geoffrey E. Hinton, and Ronald J. Williams. Learning internal representations by error propagation. In David E. Rumelhart and James L. Mcclelland, editors, Parallel Distributed Processing: Explorations in the Microstructure of Cognition, Volume 1: Foundations, pages 318–362. MIT Press, Cambridge, MA, 1986.

Cynthia Sandor, Wanbo Li, Wouter Coppieters, Tom Druet, Carole Charlier, and Michel Georges. Genetic variants in rec8, rnf212, and prdm9 influence male recombination in cattle. PLoS genetics, 8(7), 2012.

Stephan Schiffels and Richard Durbin. Inferring human population size and separation history from multiple genome sequences. Nature genetics, 46(8):919, 2014.

Daniel R Schrider, Alexander G Shanku, and Andrew D Kern. Effects of linked selective sweeps on demographic inference and model selection. Genetics, 204(3):1207–1223, 2016.

Daniel R Schrider, Julien Ayroles, Daniel R Matute, and Andrew D Kern. Supervised machine learning reveals introgressed loci in the genomes of drosophila simulans and d. sechellia. PLoS genetics, 14(4):e1007341, 2018.

Sara Sheehan and Yun S Song. Deep learning for population genetic inference. PLoS computational biology, 12(3): e1004845, 2016.

Sara Sheehan, Kelley Harris, and Yun S Song. Estimating variable effective population sizes from multiple genomes: a sequentially markov conditional sampling distribution approach. Genetics, 194(3):647–662, 2013.

Karen Simonyan and Andrew Zisserman. Very deep convolutional networks for large-scale image recognition. arXiv preprint arXiv:1409.1556, 2014.

Josef Sivic and Andrew Zisserman. Video google: A text retrieval approach to object matching in videos. In null, page 1470. IEEE, 2003.

Chris CR Smith and Samuel M Flaxman. Leveraging whole genome sequencing data for demographic inference with approximate bayesian computation. Molecular ecology resources, 2019.

Jeffrey P Spence, Matthias Steinrücken, Jonathan Terhorst, and Yun S Song. Inference of population history using coalescent hmms: review and outlook. Current opinion in genetics & development, 53:70–76, 2018.

Matthias Steinrücken, Jack Kamm, Jeffrey P Spence, and Yun S Song. Inference of complex population histories using whole-genome sequences from multiple populations. Proceedings of the National Academy of Sciences, 116(34): 17115–17120, 2019.

Lauren Alpert Sugden, Elizabeth G Atkinson, Annie P Fischer, Stephen Rong, Brenna M Henn, and Sohini Ramachandran. Localization of adaptive variants in human genomes using averaged one-dependence estimation. Nature communications, 9(1):703, 2018.

Christian Szegedy, Sergey Ioffe, Vincent Vanhoucke, and Alexander A Alemi. Inception-v4, inception-resnet and the impact of residual connections on learning. In Thirty-First AAAI Conference on Artificial Intelligence, 2017.

Jonathan Terhorst, John A Kamm, and Yun S Song. Robust and scalable inference of population history from hundreds of unphased whole genomes. Nature genetics, 49(2):303, 2017.

Luis Torada, Lucrezia Lorenzon, Alice Beddis, Ulas Isildak, Linda Pattini, Sara Mathieson, and Matteo Fumagalli. Imagene: a convolutional neural network to quantify natural selection from genomic data. BMC bioinformatics, 20 (9):337, 2019.

Rémi Tournebize, Valérie Poncet, Mattias Jakobsson, Yves Vigouroux, and Stéphanie Manel. Mcswan: A joint site frequency spectrum method to detect and date selective sweeps across multiple population genomes. Molecular ecology resources, 19(1):283–295, 2019.

Fernando A Villanea and Joshua G Schraiber. Multiple episodes of interbreeding between neanderthal and modern humans. Nature ecology & evolution, 3(1):39–44, 2019.

Ke Wang, Iain Mathieson, Jared O’Connell, and Stephan Schiffels. Tracking human population structure through time from whole genome sequences. bioRxiv, page 585265, 2019.

Alexander T Xue, Daniel R Schrider, Andrew D Kern, Ag1000G Consortium, et al. Discovery of ongoing selective sweeps within anopheles mosquito populations using deep learning. bioRxiv, page 589069, 2019.

Burak Yelmen, Aurélien Decelle, Linda Ongaro, Davide Marnetto, Corentin Tallec, Francesco Montinaro, Cyril Furtlehner, Luca Pagani, and Flora Jay. Creating artificial human genomes using generative models. bioRxiv, page 769091, 2019.

Manzil Zaheer, Satwik Kottur, Siamak Ravanbakhsh, Barnabas Poczos, Ruslan R Salakhutdinov, and Alexander J Smola. Deep sets. In Advances in neural information processing systems, pages 3391–3401, 2017.

Melinda A Zeder. Domestication and early agriculture in the mediterranean basin: Origins, diffusion, and impact. Proceedings of the national Academy of Sciences, 105(33):11597–11604, 2008.

